# Brain injury reactivates a developmental program driving genesis and integration of transient LGE-class interneurons

**DOI:** 10.1101/2025.09.30.679475

**Authors:** G. Nato, M. Fogli, N. Marichal, V. Cerrato, G. Turrini, V. Proserpio, I. Ghia, G. Zanotto, I. Molineris, M. Bergami, S. Oliviero, P. Peretto, B. Berninger, A. Buffo, F. Luzzati

## Abstract

Brain lesions can unlock a latent neurogenic potential in parenchymal astrocytes. However, the identity of their neuronal progeny has remained unclear. Here, we show that neurons generated by striatal astrocytes following excitotoxic lesions are transient, yet they reach advanced stages of morphological and functional maturation and integrate into cortico-striatal-thalamic circuits. Single-cell RNA-seq mapping onto an embryonic reference revealed that these cells are not related to adult striatal neuron types but instead belong to the LGE-MEIS2/PAX6 interneuron class. Notch abrogation, which mimics neurogenic activation, drives both cortical and striatal astrocytes toward this same interneuron class, revealing a shared intrinsic commitment. In primates, LGE-MEIS2/PAX6 cells transiently populate the embryonic striatum and cortex, and through spatial transcriptomics, we reveal that in mice these cells are also present and widely distributed throughout the telencephalon during embryonic and postnatal development. Thus, unlike other vertebrates in which adult telencephalic astroglia preserve the potential to generate constitutive region-specific neurons, the homologous cells in mammals converge on the generation of a specific transient neuron class, possibly representing a reservoir for circuit plasticity in adult life.

## INTRODUCTION

The extraordinary neuronal diversity of the adult brain originates during development from a periventricular germinal layer subdivided into committed progenitor domains (Kriegstein and Alvarez-Buylla, 2009). In most adult mammals, astroglial-like cells derived from several telencephalic domains remain neurogenic in two specialised niches: the subgranular zone (SGZ) and the subventricular zone (SVZ) (Bond et al., 2021; Obernier and Alvarez-Buylla, 2019). Like their embryonic ancestors, SGZ progenitors in the hippocampus generate dentate gyrus granule cells. By contrast, in the SVZ, the descendants of both cortical and subcortical progenitors converge towards lateral ganglionic eminence (LGE)-class interneurons that differentiate into distinct OB interneurons subtypes (Merkle et al., 2007, 2014; Schmitz et al., 2022). Whether other neuron types can be produced in the adult brain remains unclear. Under specific conditions new neurons have been described outside the canonical niches, particularly in the striatum and to a lesser extent in the cortex (Magnusson and Frisén, 2016). In the striatum, neurogenesis occurs physiologically in pre- pubertal guinea pigs (Luzzati et al., 2014), adult rabbits (Luzzati et al., 2006), monkeys (Bédard et al., 2002) and humans (Ernst et al., 2014), and under neurodegeneration in mice and rats (Arvidsson et al., 2002; Luzzati et al., 2011a; Nato et al., 2015; Thanou et al., 2025). These neurons may derive from the SVZ (Gordon et al., 2007; Liu et al., 2009; Luzzati et al., 2011a), but are also consistently generated locally (Luzzati et al., 2006, 2011a; Magnusson et al., 2014; Nato et al., 2015), in some cases such as the rabbit caudate nucleus or the mice quinolinic acid (QA)- lesioned striatum, almost exclusively. Indeed, brain lesions can unlock a latent neurogenic potential in cortical and striatal astrocytes (Buffo et al., 2008; Fogli et al., 2024; Luzzati et al., 2014; Magnusson et al., 2014; Sirko et al., 2023). At least for striatal astrocytes the prevalence of neurogenic potential is comparable to that observed in SVZ astrocytes, potentially encompassing the entire population (Cebrian-Silla et al., 2025; Fogli et al., 2024). Despite this widespread potential and robust neurogenic responses, the damaged brain fails to regenerate, raising the question of where parenchymal neurogenesis deviates from normal development. In both the neocortex and striatum lesion-induced neurogenesis exerts beneficial roles after stroke (Liang et al., 2019),(Butti et al., 2012; Jin et al., 2010), however the identity and integration capacity of the newborn neurons is still unknown.

In the striatum, the generation of medium spiny neurons (MSN), the principal striatal cell type, was initially proposed (Arvidsson et al., 2002; Parent et al., 2002), but not subsequently confirmed (Kreuzberg et al., 2010; Liu et al., 2009; Luzzati et al., 2011a; Magnusson et al., 2014; Wei et al., 2011). Other studies reported that some striatal neuroblasts express the transcription factor Sp8 and the 5HT3A receptor, typical of multiple LGE- and CGE-class interneurons (Frazer et al., 2017; Ma et al., 2012; Vucurovic et al., 2010) which were not reported to contribute to the striatum (Gokce et al., 2016; Marin et al., 2000). However, the 5HT3A receptor is expressed by numerous MGE-derived striatal interneurons (Muñoz-Manchado et al., 2016), highlighting the need for a more comprehensive molecular profile to ascertain neuronal identity. Moreover, in all models of parenchymal neurogenesis analysed so far, in physiological or pathological conditions, most if not all newborn neurons are short-lived (Arvidsson et al., 2002; Chen et al., 2004; Luzzati et al., 2006, 2011a, 2014; Ohira et al., 2010). Whether this reflects a failure to integrate into pre-existing circuits remains to be determined. In canonical niches, newborn neurons maturation and survival critically depend on electrical activity and circuit integration, with about half of them undergoing negative selection (Lin et al., 2010; Petreanu and Alvarez-Buylla, 2002). At the same time, during development, transient neuronal populations exist that integrate only temporarily to support circuit formation and refinement (Cocas et al., 2016; Damilou et al., 2024), but whether such cells can re-emerge in adulthood is unknown.

Here, we show that QA-induced striatal neuroblasts (STR-nbl) mature and integrate into the pre-existing circuits. Their transcriptional profile does not match with any adult striatal neuron lineage but rather to a GABAergic interneuron class transiently populating multiple forebrain regions, including the striatum, during development. The adult brain parenchyma thus maintains a latent potential to generate a previously unrecognized facultative neuron type, possibly involved in circuit plasticity during development and in adult life.

## RESULTS

### STR-nbl have a transient life and retain the expression of DCX during their entire life

As previously described, QA-lesion induced a strong local neurogenic response with numerous neuroblasts dispersing in the ventro-medial caudoputamen (Fogli et al., 2024) (<u>SupplementaryVideo1</u>). To unravel the identity and integration capacity of these neuroblasts, we first assessed their survival and maturation by administering BrdU at 4 wpl, when neurogenesis is already active(Nato et al., 2015) (Fig.1A). At 14d post-BrdU injection (n=6) numerous BrdU+/DCX+ cells were present in the striatum but their number dropped by about 80% at 35d (n=5; Fig.1B,E, Fig.S1A,B). By contrast the number of BrdU+/DCX- cells remained stable (Fig.1B,E; SupplementaryTable1), indicating a strong loss of newly generated cells between the two time points. BrdU+ cells never expressed DARPP-32, a MSN marker, at any time points, while at both 14d and 35d a similarly low number expressed NeuN, a marker of neuronal maturity and the interneuron markers nNOS, and Calretinin (Fig.1C, Fig.S1C,D; SupplementaryTable1). These cells however were still largely expressing DCX at 35d (69±3% of the cells; Fig.1D, Fig.S1D). In conclusion, as in other models of striatal neurogenesis, the great majority of newborn neuroblasts live transiently and do not express characteristic striatal neuron markers while retaining DCX expression for their entire life.

**Figure 1.**
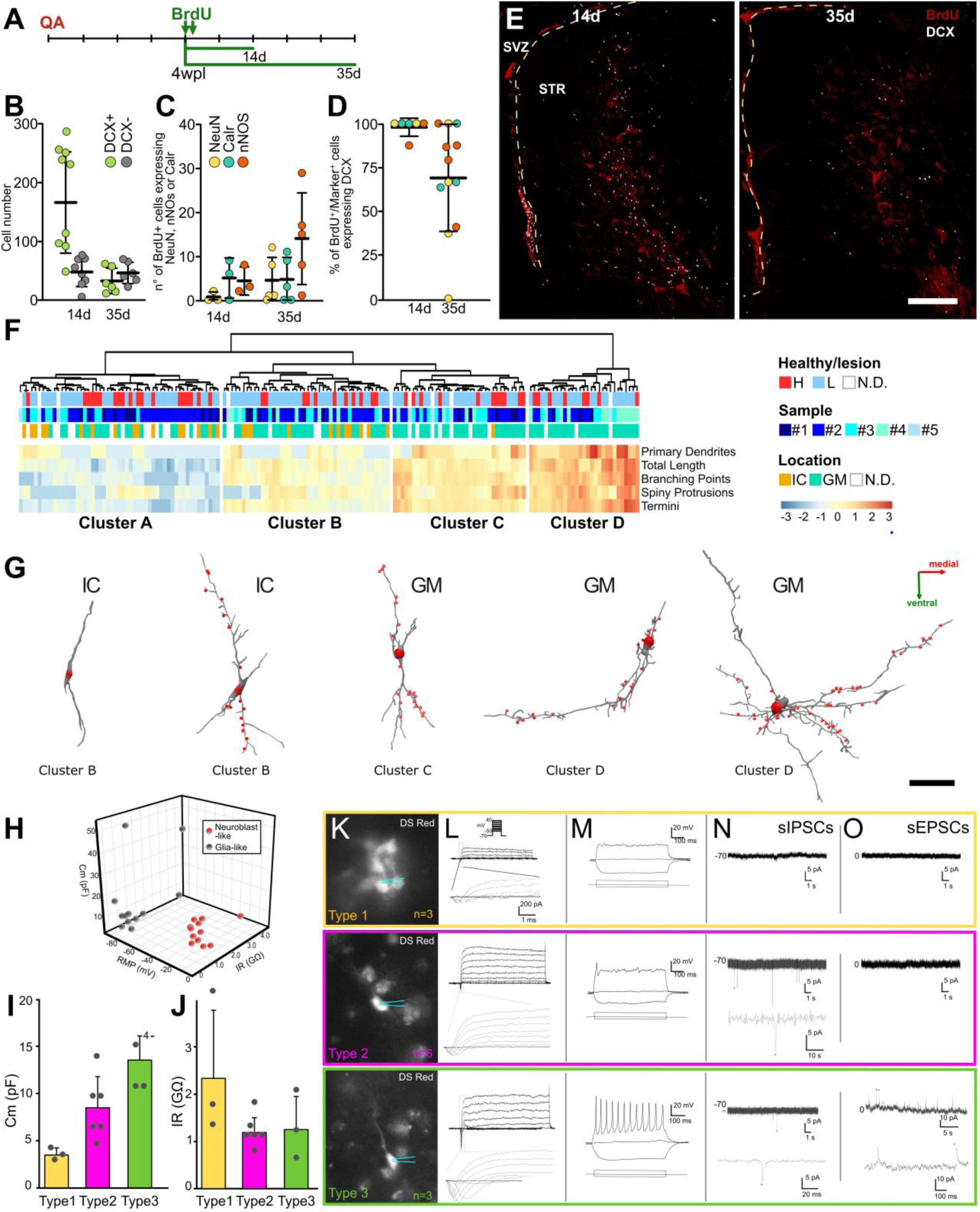
Survival and maturation. **A)** Experimental design. **B)** Number of BrdU+/DCX+ and BrdU+/DCX- cells in 2 50 μm-thick slices per animal. **C)** Number of BrdU+ cells expressing NeuN, Calretinin, or nNOS at 14d and 35d post BrdU injection. **D**) Percentage of newborn cells expressing markers of mature neurons (NeuN, Calr, nNos) retaining DCX expression (14d: n=3; 35d: n=5). **E**) Striatal neurogenic area labeled for DCX and BrdU at 14d and 35d. Note that BrdU+ but not DCX+ cells decrease with time. **F)** Hierarchical clustering based on primary dendrites, total dendritic length, branching points, spiny protrusions. **G)** 3D reconstruction of representative neurons, clusters and location in IC or GM location is indicated. **H)** 3D scatterplot showing the membrane capacitance (Cm), resting membrane potential (RMP), and input resistance (IR) of glial-like (gray) and neuroblast-like (red) cells. **I,J)** Cm and IR of Type 1, 2 and 3 neuroblasts (statistics in SupplementaryTable1). **K)** Representative fluorescence images of recorded cells. **L)** Leak subtracted currents of representative cells in voltage-clamp recordings. Type 1 cells display little outward non-inactivating currents and negligible fast inward currents. Type 2 and 3 cells display increasing amplitude of outward and especially transient inward currents.**M)** Representative firing patterns in current clamp mode after injection of depolarizing current steps. **N,O)** Examples of recorded sEPSCs (N) and sIPSCs (O). Scale bars: 200 µm in E.

### STR-nbl can attain complex morphologies and show dendritic spines

To clarify the degree of maturation reached by STR-nbl and their potential identity, we next performed high-resolution serial section 3D reconstructions (voxel size: 0.12×0.12×2µm) of 161 DCX+/RFP+ cells, 14 days after *DsRed*-encoding retrovirus injection, before the main survival drop (n=4; <u>SupplementaryVideo2</u>). Reconstructed cells displayed marked heterogeneity (Fig.1G, Fig.S1G) from the smallest and simplest cells, predominantly unipolar, with few spines, to large and complex cells, multipolar, with long branched dendrites and numerous spines (Fig.1F,G, Fig.S1G*,H). As observed in other models(Luzzati et al., 2011a, 2014; Thanou et al., 2025), part of the reconstructed neuroblasts resided in the internal capsule fiber bundles (Fig.S1E,F), often entirely included in it (IC n=23 and GM cells n=106, ratio of 1:4.5; <u>SupplementaryVideo1</u>). IC cells were generally simpler, but often more complex than typical bipolar migrating cells, and accordingly had similar spine density as GM cells (Fig.1F,G, Fig.S1H, SupplementaryTable1). Interestingly, like many OB interneurons and postnatal SVZ cells migrating to the corpus callosum (Le Magueresse et al., 2011), no STR-nbl showed distinguishable axons and most of them were clearly axonless. The STR-nbl thus acquire complex and highly heterogeneous morphologies. The presence of spines, the white matter tropism, and the lack of an axon clearly distinguish these cells from typical striatal interneurons.

### STR-nbl functionally mature and integrate into pre-existing circuits

To assess the functional maturation and connectivity of STR-nbl, we conducted whole-cell patch-clamp recordings 14 days after *DsRed*-encoding retrovirus injection, in acute slices at 5 wpl (Fig.1H-O). Labelled cells comprised both neurons and glia, which could be distinguished by their morphology and electrophysiological characteristics (Fig.1H).

In line with the variation in morphological complexity, STR-nbl displayed corresponding differences in functional maturation (Fig.1F-O). Type 1 cells (n=3) displayed the lowest Cm (Fig.1I), highest IR (Fig.1J), consistent with a smaller surface area containing less ion channels and transporters, and did not generate action potentials upon depolarizing current steps (Fig.1M). Type 2 cells (n=6) had a higher Cm, and could fire a single spikelet when depolarized (Fig.1M). Type 3 cells (n=3) showed the largest Cm and were able to fire repetitive action potentials in response to depolarizing current steps (Fig.1M), thus representing the more mature cells.

To examine lesion-induced neuroblast integration, we recorded Excitatory and Inhibitory Post Synaptic Currents (sEPSC, sIPSC) in voltage-clamp. While Type 1 cells lacked synaptic activity (Fig.1N,O), 3/6 of Type 2 neuroblasts received sEPSC at a low frequency (Freq=0.09 Hz; Fig.1N). Interestingly, 3/3 of Type 3 cells displayed sEPSC (Freq=0.06 Hz) and 2/3 also exhibited sIPSC (Freq=0.22 Hz; Fig.1N,O).

Overall, QA-lesion induced neuroblasts exhibit electrophysiological features typical of newborn neurons(Carleton et al., 2003). Despite their short lifespan, STR-nbl functionally integrate into brain circuits and preferentially receive excitatory inputs, which in the striatum typically derive from long-range afferents.

### Newborn STR-nbl receive long-range afferents

To trace the origin of synaptic inputs to STR-nbl, we used a monosynaptic retrograde tracing method based on sequential injections of modified retro and rabies viruses(Bergami et al., 2015; Deshpande et al., 2013) (Fig.2A). Retrovirus encoding RFP, rabies glycoprotein G, and TVA receptor was injected 3 weeks after QA lesion, followed 10 days later by RABV-GFP, and a 4-day chase. Double-infected cells (RFP+GFP+) were “starter cells” spreading RABV-GFP to first-order presynaptic afferents (RFP-GFP+; Fig.2A-C). To target the entire neurogenic region, we varied the injection sites across specimens (Fig.S2, Fig.2B,C; SupplementaryTable2; n=15). Only in three specimens the starter cells were exclusively DCX+ STR-nbl (Fig.2B,C,Fig.3A,Fig.S2A,B) while all others included SVZ DCX+ neuroblasts (SVZ-nbl; Fig.2D) and/or striatal DCX- cells (STR-glia; Fig.2E). Based on starter composition, specimens were divided into four groups: *Glial-only* (n=3) and *SVZ-only* (n=3) included exclusively STR-glia and SVZ-nbl, respectively. Animals containing STR-nbl were assigned to either the *STR&SVZ-Mixed* (n=7) or *STR-only* (n=8) groups, depending on the presence or absence of SVZ-nbl starter cells, respectively (Fig.2B,C, Fig.3A,B). *STR-only* specimens resulted from more lateral injections, had more STR-glia and fewer STR-nbl starter cells (Fig.3A). These STR-nbl starters were distributed more ventro-laterally and interestingly had longer and more branched dendrites (Fig.3C,D; SupplementaryTable1). Notably, only STR-nbl starters formed synaptic-like contacts with GFP+RFP– axons, often with varicosities typical of cortico-striatal inputs(Kincaid et al., 1998) (Fig.2B’’,C’’,D).

**Figure 2.**
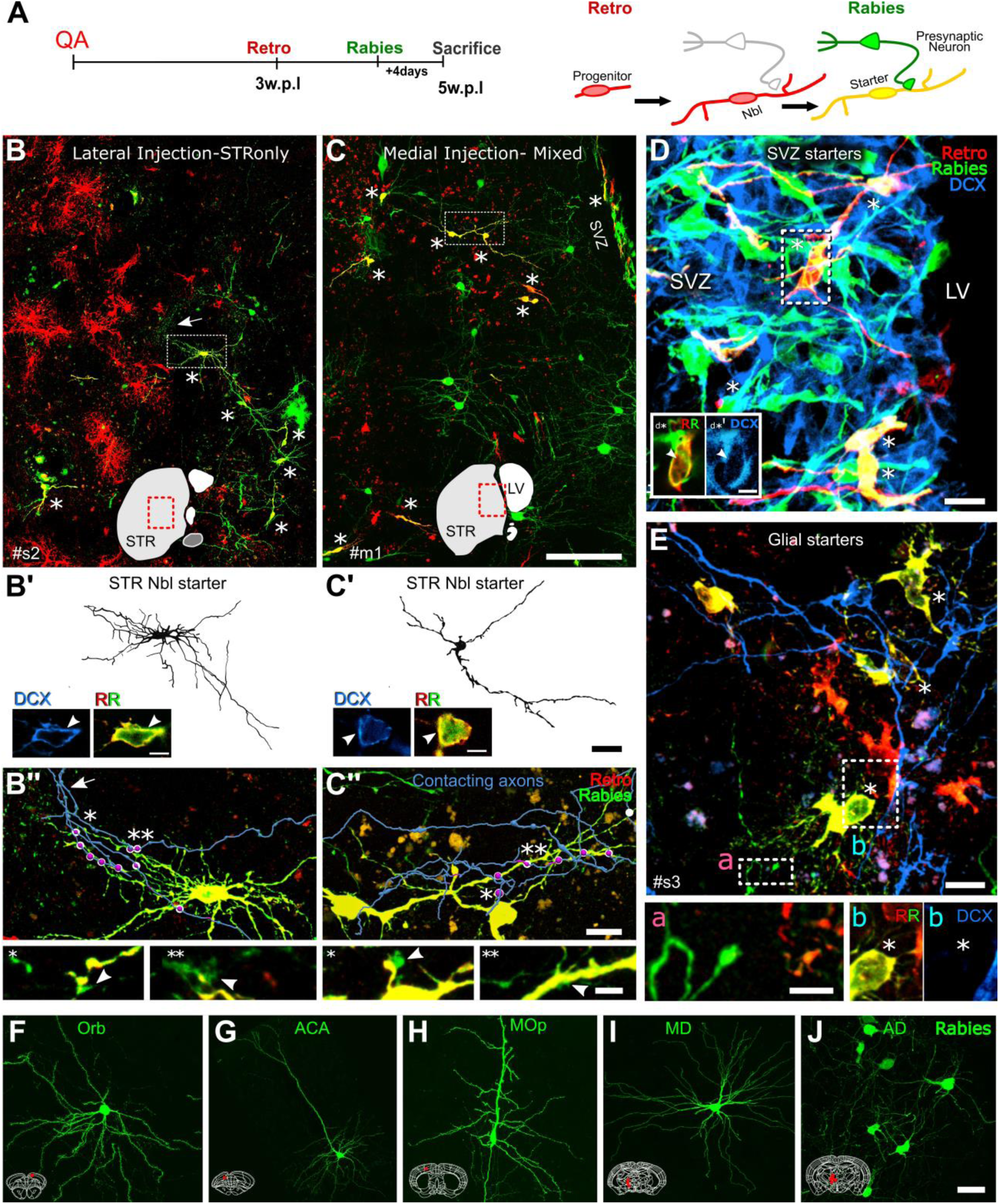
Starter cells. **A)** Experimental timeline. **B,C)** Labeling for RFP (red, retrovirus) and GFP (green, rabies virus) in the striatum of a STRonly **(B)** and a Mixed **(C**) specimen; asterisks indicate neuronal starters, arrow in B points to axonal branches. **B’,C’)** 3D reconstruction of the neuroblasts in the dotted rectangle in B and C. A single confocal plane of the cell body shows DCX expression (light blue) in these starter cells. **B’’,C’’)** Higher magnification of the dotted area in B,C. In blue segmentation of GFP+/RFP- presynaptic axons, with numerous boutons en passant, contacting the starters. Contact points resembling synaptic contacts (**\*, \*\***), are highlighted in magenta and shown at higher magnification, single planes, in the insets. **D)** DCX+ starter cells in the SVZ (*). LV: Lateral Ventricle; inset: single focal plane of a starter cell body. **E)** DCX-glial starters inside the lesioned areas (*). A GFP-positive fiber runs close to one of them without contacting it (**a**). High magnification of a starter cell (**b**). **F-J)** Examples of GFP+ presynaptic neurons located in the cortex ORB (**F**), ACA (**G**), MOp (**H**), and in thalamic MD (**I**) and AD (**J**). Scale bars: 100 µm in B,C; 20 µm in B’, C’; 10 µm in B’’,C’’,D,E; 3 µm in the inset of B’ and C’; 4 µm in B’’*,B’’**,C’’*,C’’**; 2 µm in D*,E*; 20 µm in F-J.

**Figure 3.**
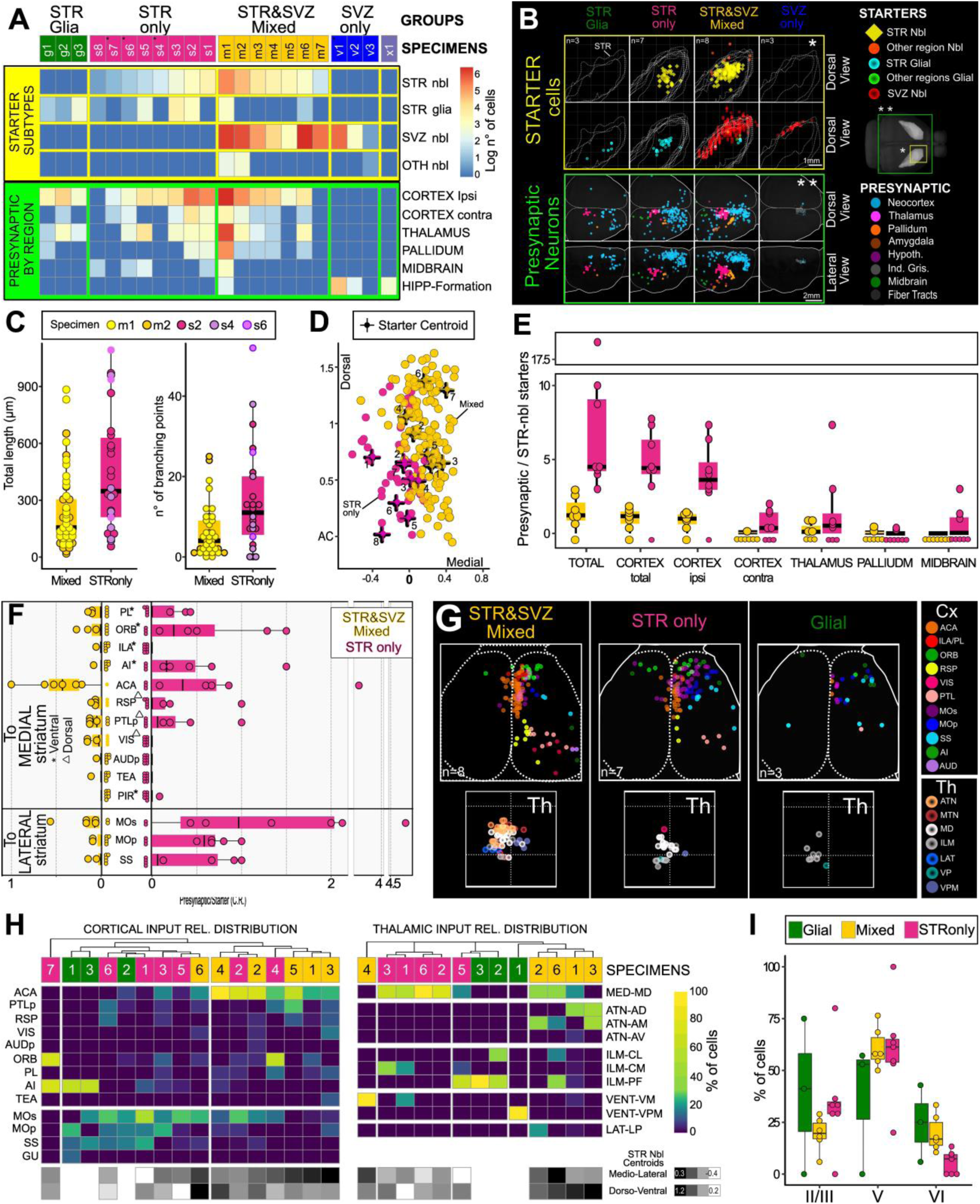
Whole-brain mapping of striatal neuroblast afferents. **A)** Heatmap of the number of main starter subtypes and presynaptic neurons in each specimen. X1 lacked starter cells in the sampled sections but had few presynaptic neurons in the indusium griseum. **B)** Spatial distribution of starter and presynaptic cells divided by group and overlapped in a common reference space. **\*)** top view of the striatum with parenchymal neuroblasts starters (first row) or glial and SVZ starters (second row). **\*\*)** dorsal (first row) and lateral (second row) views of presynaptic cells. Layer 6b cells are shown as smaller dots. **C)** Box-plot of total dendritic length and branching points of STR-nbl starters in *STR-only* and *STR&SVZ-Mixed* specimens. **D)** Front projection, aligned to the STR axis, of STR-nbl starters in different specimens. Crosses indicate the distribution centroid in each specimen. **E,F)** Box plots of the connectivity ratio in *STR-only* and *STR&SVZ-Mixed* specimens in main regions (E) and in cortical areas (F). In F, smaller dots below each bar represent specimens lacking cells in that area. Areas are grouped by preferential projection to the lateral or medial striatum, with asterisks and triangles indicating preferential targeting of the dorsal or ventral part, respectively. **G)** Top view of the cortical (Cx) and thalamic (Th) presynaptic neurons colored by cortical or thalamic subregion. Ball diameter: 250 µm. Thalamus is at higher magnification. **H)** Relative distribution (% of total) of presynaptic neurons in cortical (left) and thalamic (right) sub-regions. Specimens are indicated by numbers, colored by group and ordered by hierarchical clustering. Bottom lines indicate the position of the STR-nbl centroids. **I)** Box plots showing the relative layer distribution of cortical cells.

Presynaptic neurons were mainly in the neocortex and thalamus, the principal striatal inputs(Hunnicutt et al., 2016), with numbers also varying widely among specimens (Fig.2F-J; Fig.3A,B; Fig.S3; Fig.S4A,B). Multiple linear regression showed that STR-nbl but not SVZ-nbl or STR-glia predicted presynaptic neuron counts (Fig.S4C,D; SupplementaryTable1). Interestingly, the number of presynaptic cells per STR-nbl starter, or connectivity ratio, was higher in *STR-only* than in *STR&SVZ-Mixed* specimens (Fig.3E; SupplementaryTable1) suggesting that lateral cells may receive more inputs, possibly by virtue of their higher morphological complexity. Regression analysis confirmed this hypothesis, revealing that the centroid position of STR-nbl distribution was a strong predictor of presynaptic neuron numbers (Fig.S4E; SupplementaryTable1).

Collectively, these analyses indicate that STR-nbl starters receive long-distance glutamatergic inputs from the neocortex and thalamus that vary in strength in different striatal regions.

### Cortical and thalamic innervation of STR-nbl

Cortico-striatal and thalamo-striatal projections are organized into multiple parallel circuits that are topographically organized in the striatum(Hintiryan et al., 2016; Hunnicutt et al., 2016; Lee et al., 2020). These connections divide the striatum in medial and lateral macro-domains involved in goal-directed and habitual behaviors(Kupferschmidt et al., 2017; Yin et al., 2006). Most inputs to STR-nbl originated from regions projecting to the medial striatum, where these cells mostly reside, such as the anterior cingulate cortex (ACA, in 11/15 specimens, in which they represent the 35±20% of CX inputs) and secondary motor cortex (MOs, 11/15 specimens, 25±17% of CX inputs; Fig.3F,G). By contrast, afferents of the lateral striatum, such as the somatosensory cortex (SS, 6/15 specimens, 11±8% of CX inputs) or primary motor cortex (MOp, 7/15 specimens, 13±7% of CX inputs), were less represented. The medio-lateral position of STR-nbl in *STR&SVZ-Mixed* specimens further correlated with the expected differential representation of areas projecting to the medial and lateral striatum (Fig.3H). Like typical cortico-striatal projections, these inputs originated mainly from layer 5, and to a lesser extent from layer 2/3 (Harris and Shepherd, 2015) (Fig.3I).

Thalamic inputs were found in 10/15 specimens and were always less abundant than cortical inputs (Fig.3G,H; Fig.S3,S4). Interestingly, the two major thalamic afferents of striatal MSNs, Pf and VM nuclei (Choi et al., 2018; Hunnicutt et al., 2016; Mandelbaum et al., 2019), had limited projections to STR-nbl (in 3/10 specimens for both nuclei, 33±24% and 42±42% of thalamic inputs respectively) that rather received mostly from the MD (8/10 of the specimens, 52±28% of thalamic inputs; Fig.3H), a nucleus strongly interconnected with the prefrontal cortex with which it shares also dense projections to the medial striatum (Hunnicutt et al., 2016).

Collectively, these results indicate that STR-nbl receive afferents that are consistent with their medial position, namely the medial-cortical subnetwork (Hintiryan et al., 2016; Zingg et al., 2014) and its associated thalamic nuclei.

### Astrocyte-generated striatal and cortical neuroblasts belong to the LGE interneurons class

Our data indicate that STR-nbl are clearly distinct from canonical striatal populations, yet capable of circuit integration. To uncover their lineage commitment, we performed single-cell RNA sequencing (scRNA-seq) of PSA-NCAM+ neuroblasts (Fig.S5A) lineage-traced from adult Glast- CreERT2 positive astrocytes and isolated from QA-lesioned STR. Some cells were also isolated from the contralateral OB as a reference (Fig.4A,B). Among the sequenced cells, 53 STR and 46 OB cells expressed neuronal but not glial markers. These cells expressed markers of GABAergic neurons(*Gad1*, *Gad2*, *Slc32a1*) and immature neurons but not of mature neurons or glial cells (Fig.S5B,C). Cell cycle scoring indicated that most OB and STR-nbl were post-mitotic (99 and 91%, respectively). Notably, the STR and OB cells clustered together and had very few differentially expressed genes, suggesting they may be closely related (Fig.S5D,Supplementary Table DEG).

**Figure 4.**
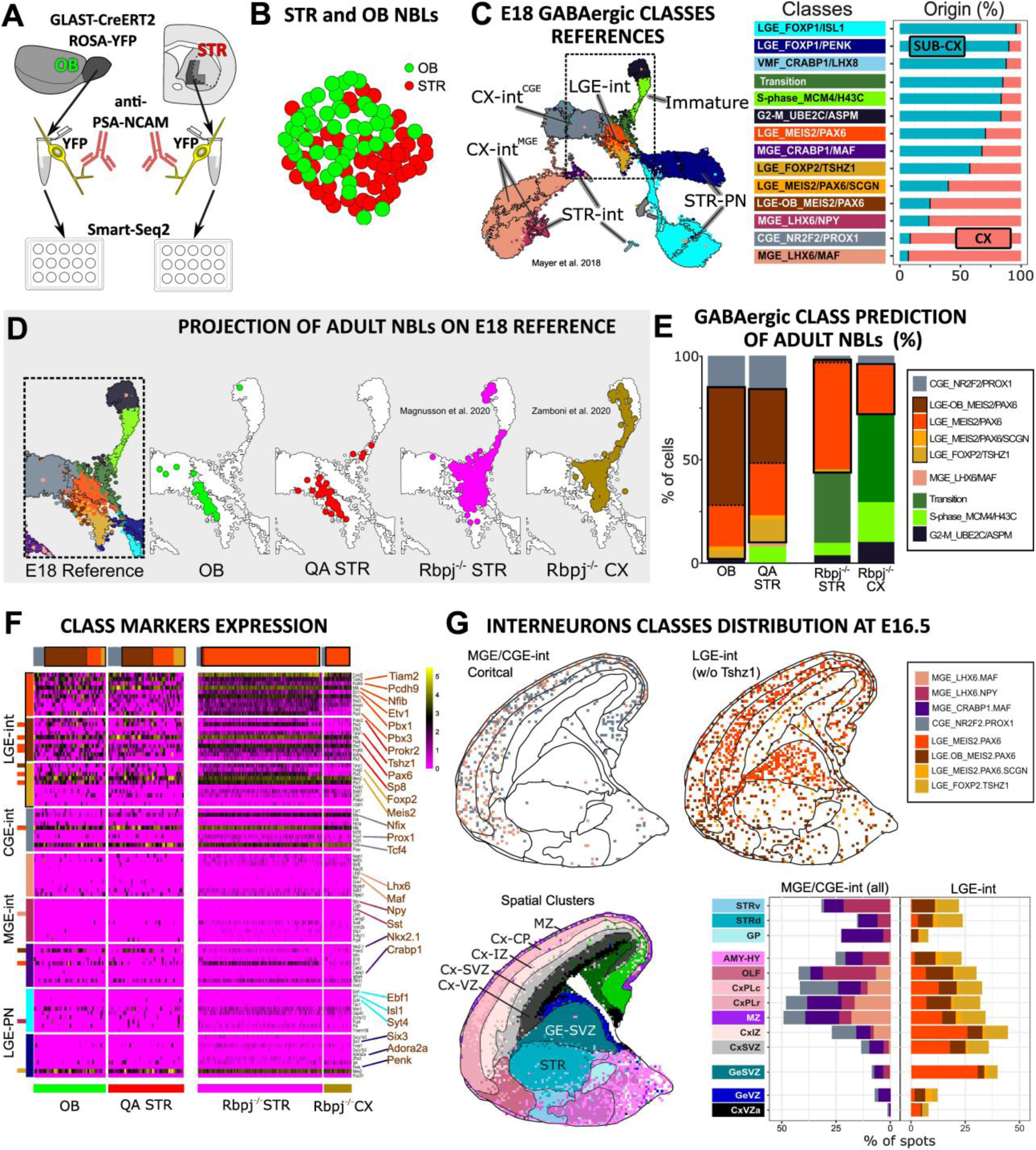
Label transfer of parenchymal neuroblasts on E18 GABAergic neurons. **A)** STR-nbl and OB-nbl isolation strategy. **B)** UMAP space showing clustering of OB and STR cells. **C)** UMAP plot of Mayers-E18 cells colored by Schmitz et al. classes. Subdivisions obtained by an independent clustering are also shown. Right panels: legend of neuron classes and fraction of reference cells in the cortex (pink) or subcortex (cyan). **D)** Magnification of Mayers-E18 UMAP corresponding to the area outlined by the black dashed rectangle in C. Projection of adult neuroblasts on the E18 reference space; samples are separated and differentially colored. **E)** Relative class predictions for each neuroblast sample. **F)** Expression of the top 10 differentially expressed genes for some E18 classes; cells are ordered according to their class attribution, as indicated by the bars on the upper side of the heatmap. Genes shared by multiple categories are indicated by lines colored as their main class. **G)** Spot distribution of MGE, CGE and LGE class interneurons (without the Tshz1 class) in an E16.5 sagittal section(Chen et al., 2022). Areal subdivision was derived from unsupervised spot clustering.

While the diversity of GABAergic neurons gradually emerges with maturation, they can already be divided into main classes at early post-mitotic stages(Mayer et al., 2018; Mi et al., 2018; Schmitz et al., 2022). To identify the class of lesion-induced STR-nbl, we projected these cells onto a large dataset encompassing all the main classes of GABAergic interneurons isolated from the mouse cortex and subcortex at E18 (Mayer-E18; Fig.4C, Fig.S5E,F) (Mayer et al., 2018). In this technique, also known as label transfer, the higher number of cells in the reference dataset results in a broader PCA space that potentially allows discriminating cell subtypes of the query datasets that could not be discriminated by clustering alone(Stuart et al., 2019). Most post-mitotic STR and OB cells mapped to LGE-class interneurons, predominantly the LGE_MEIS2/PAX6 class (Fig.4D,E) with 15%-17% assigned to CGE_NRF2/PROX1 (Fig.4E). Of note, many cells were assigned to the LGE-interneurons subclass that was defined based on adult OB immature neurons (Schmitz et al., 2022; Tepe et al., 2018), LGE-OB_MEIS2-PAX6, likely also representing more mature cells (Fig.4E, Fig.S5L). Class assignments were confirmed by the expression of class-specific markers (Fig.4F, Fig.S5G,H). Immunohistochemical analyses confirmed that striatal DCX+ neuroblasts expressed the LGE-OB_MEIS2-PAX6 marker Sp8 but not markers of other classes (Fig.S6A-F).

To more broadly analyze the cell fate potential of adult parenchymal astrocytes, we mapped onto Mayer-E18 also publicly available scRNA-seq datasets of neurons generated by cortical and striatal astrocytes after conditional Rbpj-k mutation (Magnusson et al., 2014, 2020; Zamboni et al., 2020) (Rbpj-CX, Rbpj-STR). This mutation has been proposed to mimic astrocyte neurogenic activation. Many mutant neuroblasts mapped to proliferating or early post-mitotic classes, however, the more mature cells fell into the same interneuron classes as STR and OB neuroblasts, with similar proportions (Fig.4D,E; SupplementaryTable1). None of the Rbpj-CX and few Rbpj-STR cells mapped to the adult OB subclass, consistent with a more immature state.

In conclusion, the neurogenic potential of telencephalic astrocytes converges to the production of LGE_MEIS2/PAX6 interneuron class.

### LGE-class interneurons transiently populate the embryonic and postnatal forebrain

The OB interneurons are the only adult neuron class predicted to derive from LGE-MEIS2/PAX6 neurons (Schmitz et al., 2022; Wichterle et al., 2001). Yet why parenchymal astrocytes should maintain the potential to generate OB interneurons, which do not even migrate to the OB, remains unclear. The circuit integration of striatal neuroblasts raises the possibility that, as for developmental transient neurons, their limited lifespan may be regulated by intrinsic factors. However, whether LGE_MEIS2/PAX6 class interneurons may include transient cells is still unclear. LGE-derived cells were originally reported migrating in the SVZ and intermediate zone (IZ) of the developing cortex (Anderson et al., 2001; Lee et al., 2010; Niquille et al., 2013). Putative LGE interneurons have been identified by single cell RNAseq in the developing cortical white matter and striatum of primates, including humans but their fate is unclear (Paredes et al., 2016; Sanai et al., 2011; Schmitz et al., 2022; Wang et al., 2025). By the onset of astrogliogenesis in mice (by E16.5) and humans (by GW18), putative OB interneurons are generated directly in the cortical SVZ, peaking around birth, but their fate is also unclear (Ma et al., 2012; Wang et al., 2025; Zhang et al., 2020). Interestingly, part of the Mayer-E18 LGE_MEIS2/PAX6 cells are cortical (Fig.4C, Fig.S5I), however, the precise distribution of these cells during mouse development is still unknown. We therefore mapped MGE, CGE and LGE class neurons of the Mayer-E18 cells on spatially-resolved transcriptomics data from E16.5 (Chen et al., 2022) and P7 (Joglekar et al., 2021) mouse sections (Fig.4G, Fig.S7A, Fig.5A). As expected, CGE and MGE-derived cortical interneurons distributed in CP and MZ while avoiding the striatum (Marín and Rubenstein, 2001; Villar-Cerviño et al., 2015). By contrast, LGE-MEIS2/PAX6 class interneurons were present in the cortical SVZ and IZ at E16.5 (Fig.4G, Fig.S7A), and in the corpus callosum at P7 (Fig.5A) and notably, also in the embryonic and postnatal striatum (Fig.4G, Fig.S7A, Fig.5A). To validate the presence of LGE-class interneurons in the mice postnatal striatum, we selected DEGs distinguishing LGE-class interneurons from striatal neurons (Mayer et al., 2018; Schmitz et al., 2022); (Muñoz-Manchado et al., 2018) and confirmed that Sp8 is a very specific LGE-class interneuron marker in the striatum (Fig.S5H). According to previous reports(Niquille et al., 2013), at P10, numerous Sp8+ cells were present throughout the corpus callosum (Fig.5B), but some could also be observed in the striatum (Fig.5C,H). The widespread expression of DCX at this age hampers co-labelling evaluation, however, striatal Sp8+ cells did not express the astrocyte or microglial markers SOX9 and IBA1 (Fig.S7B-C’), although a few expressed the oligodendrocyte marker SOX10 (Fig.5C). At P28, numerous DCX+/Sp8+ cells could be observed in the medial striatum (Fig.5F,I) closely resembling the STR-nbl induced by QA. Both the P10 SOX10-/Sp8+ and the P28 DCX+/Sp8+ cells almost disappeared at P90 (Fig.5C-I; SupplementaryTable1), confirming the transient presence of immature LGE-class interneurons also in the postnatal mouse striatum. Interestingly, in humans Sp8 expression have transient peaks in the prefrontal cortex and the striatum during development (Fig.S7E), supporting the presence of transient LGE-class interneurons also in humans.

**Figure 5.**
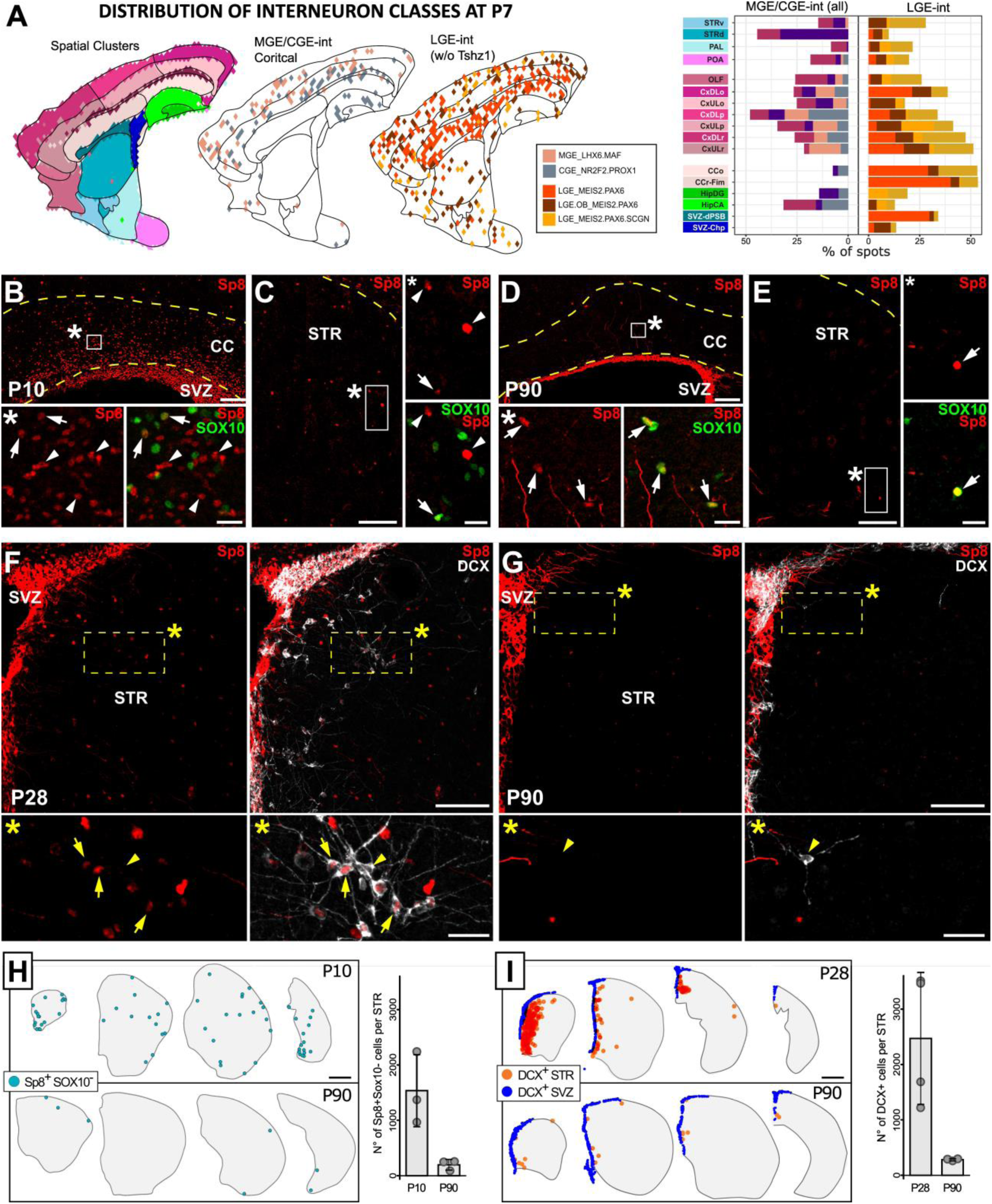
LGE-Meis2/PAX6 cells during postnatal development. **A)** Spot distribution of E18 interneuron classes at P7, as in Fig.4G. **B-E)** Sp8+ cells at the level of the CC (B,D) and STR (C,E) in P10 (B,C) and P90 (D,E) specimens. The higher magnifications (*) show that some Sp8+ cells express the oligodendrocyte marker SOX10 (arrowheads) while others do not (arrows). Dashed yellow line in B-E: corpus callosum. **F,G)** DCX/Sp8 labelled sections at P28 (F) and P90 (G). In the higher magnifications (*), both Sp8+DCX+ (arrows) and Sp8-DCX+ (arrowheads) STR-nbl can be observed. **H,I)** Representative maps of counted Sp8+Sox10- (H) and DCX+ (I) cells at different ages and striatal levels. Barplots indicate the total number of cells per striatum. Scale bars: 100 µm in B-G, 20 µm in the higher magnifications (*) of B-E, 25 µm in the higher magnifications (*) of F,G.

Overall, these analyses indicate that lesion-induced neurons in the striatum belong to a conserved population of transient interneurons populating the developing cortex and striatum. A latent potential to generate cells of this class is preserved by adult cortical and striatal astrocytes and can be activated in specific conditions.

### Heterogeneity of parenchymal astrocyte-generated neuron subtypes

MGE- and CGE-derived interneurons encompass several subtypes, partly shared across brain regions. At immature stages, transcriptomic profiles only partially distinguish these subtypes, with regional differences often being even more subtle than subtype-specific ones (Schmitz et al., 2022). The immature LGE_MEIS/PAX6 interneurons do not segregate into clearly defined transcriptional subpopulations; however, in the adult SVZ, several subtypes are known to exist that are generated from committed progenitors with distinct anatomical position and embryonic origin (Merkle et al., 2007; Obernier and Alvarez-Buylla, 2019). Interestingly, a recent comprehensive sc-RNAseq study of the adult SVZ distinguished two main lineages: dorsal and ventral (Cebrian-Silla-SVZ; (Cebrian-Silla et al., 2021) (Fig.6A). The derivatives of pallial progenitor domains are thought to generate exclusively dorsal lineage neuroblasts, while those from sub-pallial domains generate both dorsal and ventral (Cebrian-Silla et al., 2021; Merkle et al., 2007).

**Figure 6.**
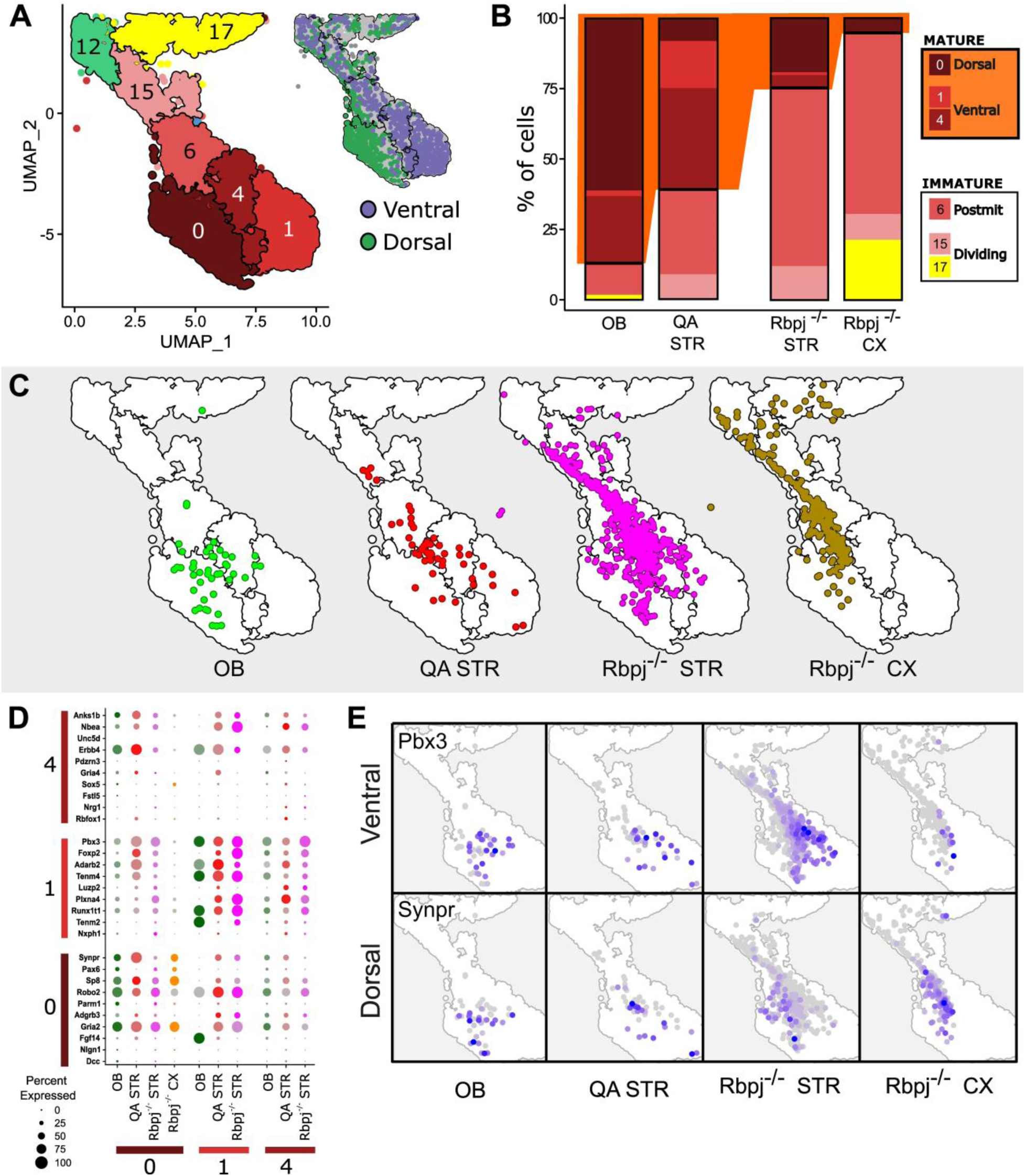
Label Transfer of parenchymal neuroblasts on adult SVZ dorsal and ventral lineages. **A)** Crop of the Cebrian-Silla cells UMAP plot showing the clusters of neuroblasts, previously defined by Cebrian-Silla et al., colored according to the original publication. In the inset, the Dorsal and Ventral lineages are shown. **B)** Relative SVZ-cluster predictions for each neuroblast sample. Note that cells from QA- lesioned striatum and Rbpj mutants mapped to more immature neuroblast clusters (15 and 6) than did OB- derived neuroblasts (SupplementaryTable1), consistent with their local origin. Notably, the ratio of neuroblasts (cluster 15) to early post-mitotic neuroblasts (cluster 6) was similar for QA-lesioned STR and RBPJs mutants, consistent with a shared early maturation program(Fogli et al., 2024; Magnusson et al., 2020; Zamboni et al., 2020). **C)** Projection of adult neuroblasts on the SVZ UMAP space; samples are separated and differentially colored. **D)** Dot plot showing the expression of the top 10 marker genes of mature neuroblast clusters obtained from (Cebrian-Silla et al., 2021). **E)** Feature plots revealing the pattern of expression on the neuroblast cells of some of the marker genes shown in D.

Cortical and striatal astrocytes share with their SVZ counterparts a distinct pallial and subpallial origin (Bayraktar et al., 2014), and we thus used label transfer to analyze whether this reflects their subtype commitment. Of note, in Rbpj mutants neuroblasts reaching more mature stages (clusters 0, 4, and 1) were markedly reduced, particularly in the cortex (Fig.6B,C; SupplementaryTable1), further supporting a reduced maturation in this model. Among neuroblasts predicted to more mature clusters (0, 4, and 1; Fig.6D), OB, Rbpj-STR and QA induced neuroblasts included both dorsal (0) and ventral (1 and 4) identities, with QA showing a bias toward ventral identities, likely reflecting the ventral location of the neurogenic response (Fig.6B, SupplementaryTable1). By contrast, mature Rbpj-CX neuroblasts were exclusively found in the dorsal cluster (0; Fig.6B,C,E; SupplementaryTable1). As with other interneuron classes, the LGE_MEIS2/PAX6 class interneurons can thus be subdivided into different subtypes that are partly shared across brain regions. In adult astrocytes, this subtype diversity correlates with embryonic origin.

## DISCUSSION

Here, we demonstrate that cortical and striatal astrocytes share with their SVZ counterparts the potential to generate LGE_MEIS/PAX6-class interneurons. In addition to OB interneurons, during development this class includes neurons that transiently populate several brain regions including the striatum. These cells can thus be viewed as a facultative striatal cell type that is generated only in specific conditions. Following injury, these cells re-emerge, mature and transiently integrate into pre-existing circuits.

### Neuroblast Maturation and Heterogeneity

STR-nbl at approximately 14d of age showed considerable variability in shapes and morphological complexity. This heterogeneity could be at least partly explained by differences in the level of maturation, as supported also by the analysis of their functional development. Indeed, simpler Type1 and Type2 STR-nbl displayed features typical of early and intermediate maturation stages, particularly of SVZ neuroblasts, including reduced membrane capacitance and increased input resistance (Carleton et al., 2003). However, Type 2 cells further showed fast inward currents and spikelet firing, indicating the presence of voltage-gated Na⁺ channels, while Type 3 STR-nbl were capable of firing repetitive action potentials. Of note, these latter features of Type2 and Type3 cells are developed relatively later in SVZ neuroblasts, only after 2 and 3 weeks, respectively. STR-nbl faster maturation may result from a shorter migratory phase, associated with the local genesis, or environmental factors such as neural activity, a well known driver of neuronal maturation (Lin et al., 2010; Petreanu and Alvarez-Buylla, 2002; Wallace and Pollen, 2024). The QA-induced neuroblasts in the ventral and lateral striatum, where resident neurons were largely depleted, displayed longer dendrites and received more synaptic inputs, possibly reflecting a competitive advantage that fosters maturation and integration. Such maturation-promoting factors could be specifically induced by QA lesion, thus explaining the lower maturation stage observed in neuroblasts derived from Rbpj mutant astrocytes, which are generated under conditions that do not spontaneously promote neurogenesis. At the same time, transcriptional analyses revealed the presence of subtypes among striatal LGE interneurons that, as for newborn OB interneurons, could be regulated by intrinsic factors (Cebrian-Silla et al., 2021). Thus, both intrinsic and local contextual factors may shape striatal LGE interneurons differentiation; however, their relative contribution remains to be defined.

### Neuroblast Fate

During both adult and embryonic neurogenesis, electrical activity promotes not only the maturation of immature neurons, but also their survival (Lin et al. 2010; Veyrac et al. 2013; Kelsch et al. 2009; Bergami et al. 2015; Priya et al. 2018; Duan et al. 2020). In line with the neurotrophic theory (Levi-Montalcini, 1987), the physiologic loss of newly generated neurons, reaching up to 50% in adult-born neurons and developing cortical interneurons, is commonly interpreted as a selection process, in which the newcomers compete for integration (Kempermann et al. 2004; Denoth-Lippuner and Jessberger 2021). However, a recent longitudinal study in the periglomerular layer of the OB showed that both surviving and subsequently eliminated neurons exhibited comparable responses to external stimuli(Su et al., 2023). Similarly, pregnancy-induced OB interneurons integrate into circuits and respond to external stimuli, yet are almost all lost after weaning (Chaker et al., 2023). As in these models, the precocious death of QA-induced STR-nbl cannot be simply explained by failed integration or lack of activity. Notably, a transient life of newborn neurons is a consistent feature across all models of adult parenchymal neurogenesis analyzed so far, in both healthy and lesioned conditions (Arvidsson et al., 2002; Chen et al., 2004; Gould et al., 2001; Luzzati et al., 2006, 2011a, 2014; Ohira et al., 2010). The environment may not be the limiting factor, as transplanted embryonic striatal precursors can survive and differentiate in both healthy and lesioned striatum (Luzzati et al., 2011a). These observations strongly suggest that the transient life of adult parenchymal neurons is governed by intrinsic factors, as proposed for other cells during development (Elorriaga et al., 2023; Southwell et al., 2012). But why might some neurons be fated to die? Before becoming fully engaged in neural signal processing, developing neurons play several structural and supportive roles that are essential for the maturation and organization of the nervous system (Cossart and Garel, 2022). These roles span all stages of circuit assembly, from proliferation to axon guidance, synapse formation and remodeling (Cossart and Garel, 2022), and can also target already established adult circuits, as observed in adult neurogenesis (Sailor et al., 2017) or embryonic neuron transplantation (Dehorter et al., 2017). In the words of Cossart and Garel (Cossart and Garel, 2022), these developmental players are then “recycled” to process information in mature circuits. From this perspective, cell selection may function as a mechanism to balance the number of neurons needed for these transient roles with those required for long-term functions. Accordingly, during development, some neuronal types undergo little to no selection, such as the cortical pyramidal neurons or early postnatal DG granule cells (Gao et al., 2014; Sarma et al., 2011). Others are only partially selected, such as the GABAergic interneurons, while others are specialized for transient functions, such as the Cajal-Retzius or Subplate cells (Elorriaga et al. 2023; Molnár et al. 2020) or the novel non-OB LGE_MEIS2/PAX6 subtypes that we described here. Cajal-Retzius and Subplate neurons also establish both local and long-range connections but ultimately die at the end of development. At least for the Cajal-Retzius cells, electrical activity promotes their death, possibly linking their fate to the completion of network maturation (Elorriaga et al., 2023; Riva et al., 2019).

### The LGE-class interneurons include populations of transient neurons during development

Early studies demonstrated the tangential migration of interneurons from the LGE to the cortical SVZ and IZ by around E14.5 in mice (Anderson et al. 2001; Pedraza et al. 2014). By the onset of gliogenesis at E16.5, OB-like interneurons begin to be generated locally within the cortex (Zhang et al. 2020; Lee et al. 2024). More recently, scRNA-seq studies identified LGE_Meis2/PAX6 cells in the macaque cortex (Schmitz et al., 2022) and closely related cells were also described in humans (Wang et al., 2025), referred to as dorsal LGE immature neurons (dLGE-int). Immunohistochemical analysis in macaques (Schmitz et al., 2022) proposed that LGE_MEIS2/PAX6 cells may populate the arc (Kim et al., 2025; Paredes et al., 2016), a prominent accumulation of immature neuroblasts along the Muratoff bundle, a fibre tract carrying numerous cortico-striatal fibers (Bullock et al., 2022). From this region, that is close to the dorsolateral corner of the LV, they disperse into the surrounding white matter, extending up to the subplate. This population may also correspond to the SVZ-derived neuroblasts associated with CC fibers connected with the prefrontal cortex in postnatal humans, rabbits and mice (Le Magueresse et al., 2011; Luzzati et al., 2003, 2006; Sanai et al., 2011). Supporting this interpretation, a recent MERFISH analysis in humans identified dLGE-int cells not only in the cortical white matter but also in the subplate and layer 6b (Wang et al., 2025). Here, we supported and extended these findings by providing the first comprehensive description of LGE_MEIS2/PAX6 neuron distribution across the embryonic and postnatal mouse telencephalon, using whole-transcriptome spatial transcriptomics. In the cortex, as in primates, these cells preferentially settled in the IZ/CC and in the subplate/layer 6. Interestingly, consistent with previous scRNA-seq data in macaque embryos (Schmitz et al., 2022), we also detected LGE_MEIS2/PAX6 cells in the mouse striatum during development. Immunohistochemical analysis of Sp8, a specific marker of these cells in the striatum, showed that at weaning, they are particularly enriched in the rostro-medial striatum.

Bioinformatic reconstruction of initial class differentiation fate in mice clearly indicates that the only adult neuron types belonging to the LGE_Meis2/PAX6 class are the OB interneurons (Schmitz et al., 2022). This conclusion is also supported by *in utero* transplantation of early LGE cells (Wichterle et al., 2001) and BrdU injections at P1 (Ma et al., 2012), when the production of these cells peaks in the cortex (Lee et al., 2024; Zhang et al., 2020). This implies that most, if not all, the LGE_MEIS2/PAX6 class neuroblasts in the embryonic and postnatal cortex and striatum represent a novel and conserved family of transient neurons. This interpretation is further supported by their strong tropism for white matter tracts and the subplate, regions rich in transient cells, both GABAergic and Glutamatergic, during development (Molnár et al., 2020; Niquille et al., 2013; Qu et al., 2016). Of note, our analyses suggest that this neuron class may also transiently populate the LOT, where, unlike the CC, only transient glutamatergic neurons have been demonstrated so far(Ruiz-Reig et al., 2017; Tomioka et al., 2000).

Like these pallial embryonic populations of transient neurons, in the QA lesioned striatum, under progressive degeneration (Luzzati et al., 2011a) or in the postnatal guinea pig(Luzzati et al., 2014), newborn STR-nbl showed a strong tropism for IC fiber bundles and adjoining gray matter. In addition, unlike adult-generated neuroblasts in canonical niches, that initially integrate into local circuits(Ming and Song, 2011; Panzanelli et al., 2009), STR-nbl received mainly glutamatergic inputs originating from long-range cortical and thalamic afferents, whose projections run through the internal capsule. STR-nbl may thus represent an actual striatal cell type, that however, in contrast to the classic ones, is characterised by a transient life and is generated on demand only in specific conditions, such as during embryonic and postnatal development, or after lesion. Like other transient cell types during development, LGE_MEIS2/PAX6 neuroblasts may be endowed with some plastic roles likely related to the formation and/or refinement of striatal circuits. The roles played by embryonic transient cells are still largely unclear. In white matter tracts, they can act as guideposts, attract growing axons and support their fasciculation (Niquille et al., 2013), while in the subplate, they establish intermediary connections with thalamic fibers that are thought to be important in proper wiring of the neocortex (Molnár et al., 2020). To what extent some of these roles are played by the LGE-interneurons remains to be clarified. Interestingly, across different models of striatal neurogenesis, the topographical distribution of LGE-MEIS2/PAX6 neuroblasts varies greatly. In mice, it depends on the lesion site(Fogli et al., 2024; Luzzati et al., 2011a), they concentrate dorsomedially in both young mice and adult rabbits, while they are abundant ventrolaterally in guinea pigs. Like in the cortex, the basic unit of cortico-striato-thalamic circuits is relatively similar across the striatum but differs functionally based on the nature of the inputs. The transient LGE_Meis2/PAX6 cells can thus interact with specific circuits during development in a species- and context-specific manner.

Brain lesions stimulate plasticity of both neurons and glial cells to support the rearrangement of surviving neuron connections, often re-awakening developmental mechanisms (Cramer and Chopp, 2000; Jones and Adkins, 2015). The striatal lesion-induced neurogenesis may thus act in concert with this remodelling attempt. Interestingly, after QA, neurogenesis starts two weeks after the lesion, in a putative recovery phase (Fogli et al., 2024). Most neuroblasts concentrate around the lesion border, where they are generated and where surviving neurons are likely undergoing a higher level of remodelling (Jones and Adkins, 2015). At least after stroke in the cortex, immature neuroblasts were shown to support recovery of connectivity and motor function (Liang et al., 2019). However, while development is completed in a few weeks, the striatal neurogenic response lasts for at least six months (Nato et al., 2015; Thored et al., 2006), with the continuous production of new transient neurons, suggesting that they may either fail to conclude their action on the circuits, or, as for canonical neurogenic niches that they may be able to support a more constitutive role in adult circuits. For instance, in the rabbit brain (Ernst et al., 2014; Luzzati et al., 2006), and possibly also in humans, adult striatal neurogenesis continues throughout life.

### Concluding remarks

As in other systems, brain lesions can trigger the activation of local stem cells and the re-emergence of developmental cell types (Aztekin, 2024; Tanaka and Reddien, 2011). At least in the cortex, the fate switch of radial glial progenitors from glutamatergic neurons to LGE_class interneurons occurs at the beginning of astrogliogenesis (Fuentealba et al., 2015; Lee et al., 2024; Zhang et al., 2020), and new astrocytes may thus keep this genetic program accessible throughout life and reactivate it on demand. In non-mammalian vertebrates, adult astroglial cells maintain not only the anatomical parcellation of the embryonic progenitors, as in mammals (Bayraktar et al., 2014), but also their radial glial form and at least part of their neurogenic commitment, thereby supporting lifelong neuronal addition and regeneration (Jurisch-Yaksi et al., 2020; Kirkham et al., 2014; Scott and Lois, 2007). Of note, in fish, OB-like interneurons are present in a region ventral to the striatum proper (Tibi et al., 2023) and are constitutively generated during adulthood (Rothenaigner et al., 2011), suggesting that this interneuron class may extend beyond the OB also in other species. The progenitors of these cells however are distinct from those of the striatal and pallial domains. The convergence of astroglial neurogenic potential from both pallial and subpallial origins onto transient LGE-MEIS2/PAX6-class interneurons may thus represent a specific mammalian adaptation. However, if and where transient LGE-class interneurons are generated during development or in response to lesion in non-mammalian vertebrates remains to be established.

The functions and dysfunctions of this emerging population of transient neurons, during development and in adulthood, await to be clarified. These cells could be harnessed for brain repair, either by exploiting their plastic potential or by reprogramming their progenitors towards earlier developmental programs.

## ACKNOWLEDGMENT

We thank Alessia Fanasca for preliminary work. This research was supported by: National Center CN3 PNRR MUR to S.O., European Union’s Horizon 2020 research and innovation programme under grant agreement no. 874758 to A.B., funds of the University of Turin and Compagnia di San Paolo (S1618 grant) to A.B and F.L., funding from the European Research Council (ERC) under the European Union’s Horizon 2020 Research and Innovation Programme (grant agreement no. 101021560, IMAGINE) and the German Research Foundation (BE 4182/11-1, project no. 357058359) to B.B. This study was also supported by MIUR project “Dipartimenti di Eccellenza” 2018–2022 and 2023–2027 to Dept. of Neuroscience “Rita Levi Montalcini”.

## AUTHOR CONTRIBUTIONS

Conceptualization, G.N., M.F., and F.L.; Investigation, G.N., M.F. and F.L.; Electrophysiology, N.M.; scRNA-seq, G.N., I.M., V.P, I.M. and S.O.; Formal Analysis, G.N., M.F., N.M., G.Z. and F.L.; Formal Analyses - Transcriptomics, V.C., G.T and F.L.; Formal Analyses – Morphology, I.G., G.N. and F.L.; Software - 3D reconstructions, F.L.; Data Analyses - Connectivity F.L; Connectivity reagents B.B and M.B. Visualization, G.N., M.F., N.M. and F.L.; Writing - Original Draft, G.N., M.F. and F.L.; Writing - Review & Editing, G.N., M.F., N.M., S.O., P.P., B.B., A.B. and F.L.; Supervision, P.P., B.B., A.B. and F.L.; Funding and resources acquisition, M.B., S.O., P.P., B.B., A.B. and F.L.

## DATA AND SOFTWARE AVAILABILITY

The original single cell RNAseq datasets have been deposited on GEO: GSE303816. All the original scripts reproducing the analyses on RNAseq as well as ImageJ and Blender scripts used for the 3D reconstructions and connectivity analysis will be made publicly available upon publication of the manuscript. The original .swc files of the morphological neuron reconstructions will be uploaded on Neuronmorpho.org.

## MATERIALS AND METHODS

### Animal procedures

The experimental plan was designed according to the guidelines of the NIH, the European Communities Council (2010/63/EU), and the Italian Law for Care and Use of Experimental Animals (DL26/2014). It was also approved by the Italian Ministry of Health (authorization 327/2020-PR) and the Bioethical Committee of the University of Turin. The study was conducted according to the Arrive guidelines.

### Mouse lines

Experiments were performed on 8-12 weeks old C57BL/6J mice and 8-12 weeks old GLAST- CreERT2 mice(Mori et al., 2006) crossed with R26R-YFP mice(Srinivas et al., 2001) to produce Glast CreERT2/R26R-YFP heterozygous for GLAST-CreERT2 and homozygous for R26R-YFP.

### Stereotaxic injections

Mice were anesthetized with a solution containing ketamine (133 mg/kg; Ketavet 100; Gellini, Aprilia LT, Italy) and xylazine (6.7 mg/kg; Rompun 20 mg/mL; Bayer, Wuppertal, Germany), positioned in a stereotaxic apparatus, and injected into the right hemisphere using a picopump (Picospritzer II, General Valve Corporation). *QA injection*: 1 µl of quinolinic acid (QA) diluted to 120 mM in 0.1 M PB was injected +0.1 mm AP, −2.1 mm ML, and −2.6 mm DV. *Retroviral injections for morphological analysis and electrophysiological recordings*: double injections of CAG-IRES-DsRedExpress2 or CAG(GW)-Psd95*YFP-IRES-dsRed retroviruses were performed at the following coordinates: +0.9 mm AP, −1.8 mm ML, and 3.2-3-4 mm DV, and +1.1 mm AP, −1.5 mm ML, and 3.0-3-2 mm DV.

*Viral injections for monosynaptic retrograde tracing*: Retrovirus: CAG-DsRedExpress2-2A-G- IRES2-TVA; Rabies viruses: EnvA-pseudotyped_SAD_ΔG-eGFP, EnvA-pseudotyped_SAD_ΔG- Gephyrin*GFP; see SupplementaryTable2 for injection coordinates.

### BrdU pulse labeling

Mice received two intraperitoneal injections of 5-bromo-2-deoxyuridine (BrdU, Sigma-Aldrich; 50 mg/kg in 0.1 M Tris pH 7.4) at 4 weeks after QA injection, 6 hours apart, and were sacrificed either 15 days (15d) or 35 days (35d) after the last injection.

### Histology

Animals were anesthetized with a ketamine (266 mg/kg) and xylazine (20 mg/kg) solution and perfused with a solution of 4% paraformaldehyde (PFA, Sigma-Aldrich) and 2% picric acid (AnalytiCals) in 0.1 M sodium phosphate buffer (PB) pH 7.4. Brains were then post-fixed for 5 hours, cryoprotected in 30% sucrose (Sigma-Aldrich) in 0.1M PB pH 7.4, embedded at −80°C in Killik/OCT (Bio-Optica), and cryostat sectioned in series of 50 µm-thick coronal sections.

### Immunofluorescence staining and image acquisitions

Sections were incubated for 48 h at 4°C in 0.01 M PBS pH 7.4 containing 2% Triton X-100, 1:100 normal donkey serum, and primary antibodies (SupplementaryTable3). For BrdU staining, sections were pre-incubated in 2M HCl for 30 min at 37°C and then rinsed in 0.1 M borate buffer pH 8.5. Sections were incubated overnight with appropriate secondary antibodies (SupplementaryTable3) and coverslipped with antifade mounting medium Mowiol (4-88 reagent, Calbiochem).

All the fluorescent images presented in this study were acquired on a Leica SP5 confocal microscope (Leica Microsystems) equipped with 40x and 63x objectives (HCX PL APO lambda blue: 40x, NA 1.25; 63x, NA 1.4). Most 40x magnification images were taken at a resolution of 0.38 µm/pixel with optical sectioning in Z every 1.5 µm, unless otherwise specified.

### Electrophysiology

Mice were deeply anesthetized with isoflurane (Forane, Abbvie), decapitated and the brains were quickly collected into a chilled artificial cerebrospinal fluid (ACSF; Composition in mM: NaCl, 85; Sucrose, 73; KCl, 2.5; NaHCO3, 25; CaCl2, 0.5; MgCl2, 7; NaH2PO4, 1.25 and glucose, 10; pH 7.4) saturated with 95% O2 and 5% CO2 (pH 7.4). Coronal brain slices (250 µm thick) containing the striatum were prepared using a vibratome (VT1200 S, Leica) and transferred to standard ACSF (Composition in mM: NaCl, 125; KCl, 2.5; NaHCO3, 25; CaCl2, 2; MgCl2, 1; NaH2PO4, 1.25 and glucose, 12; pH 7.4). Slices were incubated in standard ACSF at 34C for 1h followed by one additional hour at room temperature (21C±2°C). For recordings, individual slices were transferred into a recording chamber mounted on the stage of an upright Zeiss microscope (Axio Imager 2, Germany). Slices were constantly perfused at the rate of 1-2 mL/min with standard ACSF maintained at RT and saturated with 95% O2 and 5% CO2.

Cells were visualized by epifluorescence and infrared DIC videomicroscopy using a 40X (0.8 numerical aperture) water immersion objective. Patch-clamp whole-cell recordings were performed in a total of 24 cells using capillaries (5-10 MΩ) pulled from borosilicate glass (BF150- 86-10, Sutter Instruments) in a horizontal puller (P-1000 Micropipette puller, Sutter Instruments) and filled with intracellular solution (Composition in mM: K-gluconate, 125; NaCl, 5; HEPES, 10; EGTA, 10; MgCl2, 2; ATP, 2; Biocytin, 5 at pH 7.4). Only cells in which an evident DsRed signal was observed within the pipet tip were considered. Voltage-clamp and current-clamp recordings were obtained using a Axopatch 200B amplifier (Molecular Devices), digitized (Digidata 1440A, Molecular Devices), and acquired and analyzed with pClamp 10 software (Molecular Devices). In voltage clamp mode, voltage steps were applied from −70 mV holding potential to levels ranging from −100 to +60 mV in 10 mV steps. To subtract leak currents, we used a P4 protocol provided by Clampex10 that allowed simultaneous storage of raw and leak subtracted data.

In current clamp mode, passive and active membrane properties were recorded by applying a series of hyperpolarizing and depolarizing current steps (2-or 10-pA steps, 500 ms). The RMP was assessed immediately after break-in by reading the voltage value in the absence of current injection (I = 0 configuration). Input resistance (Rin) was calculated from the peak of the voltage response to a −20-pA, 500-ms current step according to Ohm’s law (Vhold = −70 mV). The membrane capacitance (Cm) was obtained with the following equation: membrane time constant (τm) = Rin · Cm. τm was derived from a single exponential fitted to the voltage response to −20- pA, 500-ms current step.

### Neuroblast reconstruction for morphological analysis

Confocal microscopy serial section 3D reconstructions were performed as previously described (Fogli et al., 2024; Luzzati et al., 2011b). Briefly, confocal high-resolution images at the level of the striatum were acquired with a 63X objective (N.A. 1.4), at a resolution of 0,12 µm/pixel with optical sectioning in Z every 0.8 µm, to appreciate cellular details. The optical plane sequences of subsequent sections were aligned in TrakEM2. Cells that were entirely comprised in the reconstructed volume were traced with NeuTube (https://www.neutracing.com). During tracing, all the processes shorter than 3um were considered as spines and their traces were detached from the main branches. When possible, the border of the IC was identified based on background stain. Renderings were exported from Neutube. Unbiased hierarchical clustering analysis on the reconstructed cells was performed using the R package pheatmap. The same procedure was used to reconstruct the starter cells from selected rabies virus-injected animals (see next chapter).

### Quantification and distribution analyses of Starter cells and presynaptic neurons

To reconstruct the distribution of starter and presynaptic cells, most brains were imaged during cryostat cutting to serve as a pre-aligned template. One in three sections were labelled for antibodies against RFP and GFP, DCX,and either CTIP2 or DAPI. Whole sections were acquired with a slide scanner Axio Scan (.Z1, Zeiss) equipped with a 10x objective (Plan-Apochromat 10x, NA 0.45 M27). Images of 3 planes per section were acquired at a resolution of 0.65 μm/pixel and subsequently merged into a MAX projection. Regions of higher cell densities, mainly in the striatum, were acquired at the confocal microscope with a 40X objective (voxel size 0.38x0.38x1 µm). Images of the different modalities were registered in TrakEM2, where the starter and presynaptic cells were then manually annotated as "ball objects". Aspecific local infection along the RABV injection track, as previously reported (Beier et al. 2015), was observed in the striatum and occasionally in cortical layer 6b (data not shown). Presynaptic cells in these areas were thus excluded from further analyses. Area attributions were based on CTIP2 and DAPI staining and were further verified by alignments to the Allen Brain Atlas. To this aim, low-resolution TrakEM2 exports of these brains were imported in QuickNII (https://www.nitrc.org/projects/quicknii) (Puchades et al., 2019). Annotations were based on the Allen Brain Atlas hierarchy and nomenclature. Export from TrakEM2 and cell countings per section and per animal were obtained with customized scripts in Python and R. For 3D analysis, using customized scripts, the positions of TrakEM2 ball objects were imported into Blender3D. Surfaces of the whole brain, striatum and lateral ventricle were manually drawn in TrakEM2, exported as .obj files and imported in Blender 3.79 together with the coordinates of the more rostral and caudal ends of the CC and of the anterior commissure. These points and the surfaces were used to manually align the different brains to a common reference space. To analyze the position of the starter cells, these alignments were further refined, focusing solely on the striatum. To better understand the overlap with cortical projection domains, we downloaded maps from anterograde tracer experiments that had already been segmented and registered to the Allen Brain Atlas common reference space http://www.mouseconnectome.org/CorticalMap/page/map/5. (Hintiryan et al., 2016). The segmented sections from each individual experiment were imported into TrakEM2, overlaid onto the corresponding Allen Brain Atlas sections, interpolated, and exported as 3D objects along with selected major brain subdivisions from the Atlas. These exports were then imported into Blender and aligned to the common reference space of the injected brains. Projections from different experiments were grouped into composite 3D objects representing specific cortical areas (SSp, MOp, PTL, ORB, ACA, and PL). These objects were rendered as volumetric representations, with color intensity proportional to the local thickness of the projection The following experiments were used: ACAd: SW110323-03, ACAv: SW110321-04, MOp-ll: SW110321-01, MOp-mi: SW110517-03, MOp-mo: SW121210-01, MOp-tr: SW120118-03, MOp-ul: SW121211-02 ORBl: SW120525-01, ORBm: SW120301-02, ORBvl: SW110809-02, PLd: SW120229-04, PLv: SW110613-02, PTLp-cl: SW110906-03, PTLp-cm: SW120118-04, PTLp-r: SW110418-04, SSp-BFD-ll_f_SW110419-03, SSp-BFD-mo: SW110420-01, SSp-BFD-tr_f_SW110418-03, SSp-BFD-ul_f_SW101014-04, SSp-ll: SW120525-02, SSp-mi: SW110418-01, SSp-mo: SW110516-02, SSp-tr: SW110418-02, SSp-ul: SW110419-02

### Neuroblasts phenotyping

Quantification of the number of cells expressing BrdU, DCX (14d n=8; 35d n=6) was performed for each animal on two 50 µm-thick sections (Fig. 1B). Quantification of the number of BrdU+ cells expressing NeuN, nNOS, Calretinin, and DARPP32 (Fig. 1C-D) was performed for each animal on two 50 µm-thick sections for each marker considered (14d n=3; 35d n=5).

Quantification of the number of cells expressing LHX6, NKX2.1, FOXP2, DARPP-32, Sp8, and DCX (Fig. S5) was performed for each animal on two 50µm-thick sections for each marker considered (n=3).

### Quantification and distribution analyses of Sp8 and DCX+ cells during postnatal development

Animals at P10 (n=3), P29 (n=4), and P90 (n=3) were sectioned in 6 series of 50µm-thick sections and individual series were labelled for Sp8 and Sox10 (at P10 and P90) or for DCX and Sp8 in P29 and P90. Labelled sections have been acquired at the confocal microscope (Leica Stellaris5) with a 40x objective at a resolution of 0.28x0.28x1.5µm/px, 10 focal planes. The Sp8+ and SOX10- cells have been counted on four equally spaced sections spanning the entire striatum, while the DCX+ cells were automatically counted with Ilastik in all the sections of the series (Fig. 7H,I). To estimate the total number of cells over the entire striatum, the counted cells have been multiplied by the inverse of the sampling fraction.

### Single-cell RNA sequencing

For single-cell sequencing analysis (scRNA-seq), QA was injected in GlastCreERT2-R26R animals, and tamoxifen was administered 1 week before the lesion to select striatal astrocyte progeny (Nato et al., 2015; Fogli et al., 2024). Five weeks after lesion, mice were anesthetized with isoflurane and immediately cutted using a 0.5mm Coronal Brain Matrix (Stainless Steel, Agnthos) in PBS-glucose medium. The peri-lesion part of the striatum was dissected under a Leica Stereomicroscope, ensuring to discard the SVZ. Four biological replicates containing three QA-lesioned striata each were prepared. As a control, we also collected one OB for each animal. The lesioned striatum and OB dissected tissues were then transferred to a 15 ml falcon tube containing HibernateA (Brainbits LLC, #HA). Tissue dissociation was performed with the Miltenyi Neural Tissue Dissociation Kit (130-093-231 T) and dissociated cells were labelled with a PSA-NCAM APC antibody. This surface glycoprotein is expressed by immature and migrating neurons of both the striatum and the OB. Cells that co-expressed both YFP and APC were subsequently FACS-sorted (Cell Sorter BD FACS Aria III) and subjected to SMArtSeq2 (Picelli et al., 2014). This technique is characterized by full-length coverage, high sensitivity (i.e. it detects a high gene number per cell), and high precision in transcript-level quantification. After mRNA conversion into cDNA, individual libraries were prepared by tagmentation using a transposase enzyme. Libraries were sequenced on Illumina NextSeq 500 System. Following quality controls (performed with FastQC v0.11.2 (https://www.bioinformatics.babraham.ac.uk/projects/fastqc), sequencing reads were processed with Trim Galore! v0.5.0 (https://www.bioinformatics.babraham.ac.uk/projects/trim_galore) to perform quality and adapter trimming (parameters:–stringency 3 –q 20). Trimmed reads were next aligned to the mouse reference genome (mm10/GRCm38 Ensembl release 84) using STAR v2.7.1a51 with options:– outFilterMultimapNmax 10–outFilterMultimapScoreRange 1 –outFilterMismatchNmax 999 – outFilterMismatchNoverLmax 0.04. Gene expression levels were quantified with featureCounts v1.6.1 (https://subread.sourceforge.net/, options: -t exon -g gene_name) using the GENCODE Release M23 annotation. Multi-mapped reads were excluded from quantification. Datasets have been deposited on GEO: GSE303816.

### Quality control and clustering

Data analysis was performed in Seurat v4.2.1. Cells with less than 500 detected genes and more than 10% of mitochondrial DNA were filtered out. Two batches were merged using the default Seurat integration method based on Canonical correlation analysis (CCA) using default parameters. For integration, we used log-normalized TPM, and reduced the k.weight of the IntegrateData function to 49. After integration, we calculated PCA and clustered cells using the first 10 significant PCs as assessed by the JackStrawPlot function in Seurat. A group of microglial cells clustered separately and were thus excluded from further analyses. Fiducial markers of the main neural cell types confirmed that all the remaining cells corresponded to immature GABAergic neurons. After subsetting, we recalculated the PCA and clustered cells using significant PCs. The analysis of ***differentially expressed genes (DEGs***) was performed using FindAllMarkers function on Log-transformed TPM, using the Wilcoxon Rank Sum test, only genes with an average log2 fold change =>0.5 and expressed by at least 25% of the cluster cells were considered. ***Cell Cycle scores and NSCs developmental stage scores*** were calculated using the CellCycleScoring and AddModuleScore functions of Seurat, respectively. The gene lists defining the different NSCs developmental stages were obtained from (Kalamakis et al., 2019).

### Public Datasets

***Mayer E18 dataset,*** including E18 cortical and subcortical GABAergic cells (Mayer et al., 2018) was obtained from <u>GSE104156</u> and imported into Seurat. Cells were filtered and analyzed as described; the first 50 PCs were used for clustering and UMAP plot. Different clustering depths were considered and manually annotated. The neuron class annotations produced by Schmitz et al. (Schmitz et al., 2022) were downloaded from https://dev-inhibitory-neurons.cells.ucsc.edu. Original annotation colors were derived from the original scripts at https://github.com/mtvector/dev-and-evo-of-primate-inhibitory-neurons/tree/main/mouse, for clarity we modified the color of the LGE-OB_Meis2/PAX6 from red to brown. The ***Cebrian-Silla SVZ dataset*** (Cebrian-Silla et al., 2021) was obtained from <u>GSM5039270</u> and included the original annotations. The UMAP dimensionality reduction did not include the model, which is required for map transfer, and we thus generated a new UMAP using the first 50 PCs, as reported in the original study. ***Striatal and cortical RBPJ mutant cells*** were downloaded from <u>GSM4658401</u> and <u>GSE13984,2</u>, respectively(Magnusson et al., 2020; Zamboni et al., 2020), cells originally annotated as neuroblasts were subsetted and their neuronal identity was confirmed by expression of fiducial markers and label transfer on the Cebrian-Silla SVZ dataset.

Expression of Sp8 in Human development: SP8, probe A_24_P23546, https://human.brain-map.org/ (Hawrylycz et al., 2012) Allen Human Brain Atlas.

### Label transfer

To explore the identity of STR and OB neuroblasts obtained from this study and of striatal and cortical RBPJ mutant cells (query), we projected them on the Mayer E18 or Cebrian-Silla SVZ datasets (reference) using LabelTransfer and MapTransfer functions in Seurat. This analysis has been performed on log-normalized gene counts for both reference and query databases. Label transfer robustness was evaluated by systematically varying the number of principal components (PCs) and the k.weight parameter. Across all tested combinations, the overall repertoire of predicted identities remained stable, with no major identity classes being lost or newly introduced. Furthermore, when using more than 20 PCs, individual cell label assignments showed high consistency, suggesting that the label transfer procedure yielded robust and reproducible identity predictions. We thus ended up using the default parameters for PC 1:30 and an intermediate value for the k.weight.

### Identity transfer to the spatial transcriptomics dataset

The neurons and their progenitors from embryonic day (E) 18 mouse striatal dataset(Mayer et al., 2018; Schmitz et al., 2022) were aligned with four sagittal mouse brain sections from two E16.5 specimens analyzed with stereo-seq(Chen et al., 2022) and a postnatal day 7 (P7) sagittal section analyzed with Visium(Joglekar et al., 2021). Prior to alignment, cells belonging to the developing forebrain were subsetted based on spatial coordinates. Subsequently, the cells were normalized with the function SCTransform and clustered with FindClusters (k=0.2) from Seurat v5(Hao et al., 2024), after that, markers were calculated with FindAllMarkers and inspected to annotate the cells based on the anatomic region they occupied, which was inferred by the overexpression of characteristic markers for their most abundant populations.

To align spatial cells with the Mayer-E18 dataset, an approach based on a loss function of cosine similarity between scRNA-seq and spatial data was performed with the Python package Tangram(Biancalani et al., 2021). Clusters from the Mayer-E18 dataset, following the annotation of Schmitz et al.(Schmitz et al., 2022) were fitted on each forebrain slice with the tangram function map_cells_to_space based on the normalized expression of a set of training genes. Training genes were defined with the Scanpy function tl.rank_genes_groups and the tangram function pp.adatas(Wolf et al., 2018), this process scored each cell in the slices based on their probability of belonging to the neuron classes of the E18 dataset. The Tangram-obtained scores indicate the probability of each Schmitz GABAergic neuron class being present in each spatial spot (i.e., the probability of cells from identity *i* being present in spot *j*). The log2 score distribution for all identities displayed a prominent peak at higher values, which we interpreted as a correct attribution of clusters to their respective spots. This peak appeared at similar score values across all identities, indicating a consistent and relatively uniform strength of attribution and suggesting relatively spatially segregated distributions. Interestingly, the score distributions for cortical interneurons exhibited a left-sided tail of lower values, whereas LGE-derived interneurons showed an additional, broader peak at substantially lower values. This second peak likely reflects the high transcriptional similarities between LGE interneurons subtypes and possibly also some spatial overlap. In order to avoid confounding factors, we attributed each spot to the identity with the highest score, filtering out values falling below the local minimum between the Gaussian peaks.

### Statistical analyses

For mean comparison, we assessed whether the data were normally distributed, using the Shapiro–Wilk test, and homoscedastic, using Levene’s test. Data that fulfilled those requirements were compared with the t-test in the case of two groups or with the ANOVA in the case of three or more groups. ANOVA analyses that returned significant F values were followed by Tukey’s post hoc test. For comparison of proportions, we used the Fisher’s Exact Test, suitable to compare groups with low numbers of observations.

Simple and multiple linear regression analyses were performed to evaluate the ability of different starter cell populations to predict the number of presynaptic neurons. Model quality was assessed by comparing adjusted R-squared (Adj R²) values, which indicate the proportion of variance in presynaptic neuron numbers explained by the model, as well as the associated significance (p-values). The large variability in presynaptic and starter cell counts led to violations of linear model assumptions, which were resolved by applying a logarithmic transformation. Partial F-tests were used to statistically assess whether the inclusion of additional independent variables significantly improved the amount of explained variance (Adj R²).

Generalized linear mixed effect models were used to compare variables among groups, when we had to consider that multiple observations were obtained from individual specimens and thus the group n was not composed of independent observations (Fig.3C,D). Based on data distribution, different types of models with maximum likelihood estimation of model parameters were applied (i.e. Gaussian, Tweedie, Poisson, Negative binomial). Generalized linear mixed-effect models were run with the R packages lmer4 and glmmTMB. Likelihood-ratio tests were performed with the drop1 function of the lme4 package to obtain global statistics.

All the results of statistical tests presented in this study are reported in the SupplementaryTable1.

## SUPPLEMENTARY ITEMS TITLES AND LEGENDS

### SUPPLEMENTARY FIGURES AND TABLES

In the ’Supplemental Information’ document you will find all the supplementary figures and tables.

*Supplementary Figure 1. Lesion-induced neuron survival, maturation and morphology.*

*Supplementary Figure 2. Starter cell distribution in the striatum*.

*Supplementary Figure 3. Presynaptic cell distribution*.

*Supplementary Figure 4. Linear regression analyses of presynaptic neuron distribution.*

*Supplementary Figure 5. Adult neuroblasts sc-RNAseq clustering and E18 projection.*

*Supplementary Figure 6. Immunohistochemical phenotyping of QA-lesion induced neuroblasts*

*Supplementary Figure 7. Spatial distribution of LGE_Meis2/Pax6 class cells at E16.5 and P7*.

*Supplementary Table 1. Results of all statistical analyses presented in the study.*

*Supplementary Table 2. Stereotaxic coordinates used for monosynaptic retrograde tracing experiments*.

*Supplementary Table 3. List of primary and secondary antibodies*.

### SUPPLEMENTARY VIDEOS

***Supplementary video 1: Whole striatum 3D reconstruction of STR-nbl and IC bundles***

https://youtu.be/qMo5QAUQH7s

Animated perspective view 3D reconstruction of the whole neurogenic response obtained by registering 42 consecutive 50 μm-thick coronal sections acquired at the confocal microscope at low resolution (514 focal planes, voxel size 0.76 x 0.76 x 4.09 μm). The animation starts with the 3D model of the lateral ventricle, in blue, and a stack of 3 optical planes at the level of the anterior commissure, approximately in the center of the reconstruction. The focal planes then scroll rostrally up to the anterior limit of the striatum. These images show the DCX (white) and Ki67 immunohistochemistry with a strong myelin background stain (red), thus revealing the fiber bundles of the internal capsule. By **0:13s** the 3D rendering of DCX+ cells is progressively revealed, scrolling caudally. To limit model complexity, only higher DCX intensities were segmented into 3D objects. To appreciate also the finer and fainter processes, the volume rendering of the original fluorescence signal (normal shading) was added. At **0:17s** segmentation of neurogenic foci, obtained by the segmentation of Ki67+ clusters (see Fogli et al. 2024), is revealed. At **0:18s** the contours of the lesion, traced in TrakEM2 and based on lower magnification of reactive glia (not shown) are added. At **0:31s** the segmentation of internal capsule (IC) fiber bundles traversing the striatum is shown. At **0:39s** the camera enters the rostro-ventral part of the neurogenic response, in the healthy striatum, with numerous cells in the gray matter. Proceeding caudally, some IC bundles are traversed and ultimately, the camera enters an IC bundle and follows it until the caudal end of the striatum. At **1:24s** a sagittal reslice, 12um thick, scrolls from lateral to medial showing the presence of cells in both white and gray matter. From a closer distance, at **1:42s,** note that DCX+ cells inside IC bundles are oriented parallel to the fibers.

***Supplementary video 2: High-Resolution 3D Reconstruction of Retrovirally Labelled Striatal Neuroblasts***

https://youtu.be/yIM2J04Syds

Animated perspective view of a 3D reconstruction of the central part of a QA lesioned striatum obtained by registering 10 consecutive 50 μm-thick coronal sections acquired with the confocal microscope at high resolution (500 focal planes, 0.18x0.18x1 μm). This specimen was injected with a retroviral vector carrying a DsRed reporter 14 days before sacrifice, labelled cells were restricted to the striatum. The animation begins with a rostral view, with medial on the right. After a 180° rotation, the camera moves to a caudal perspective and zooms in on individual neuroblasts, moving ventrally.

