## Supplemental Information for "Brain injury reactivates a developmental program driving genesis and integration of transient LGE-class interneurons"

### SUPPLEMENTARY FIGURES

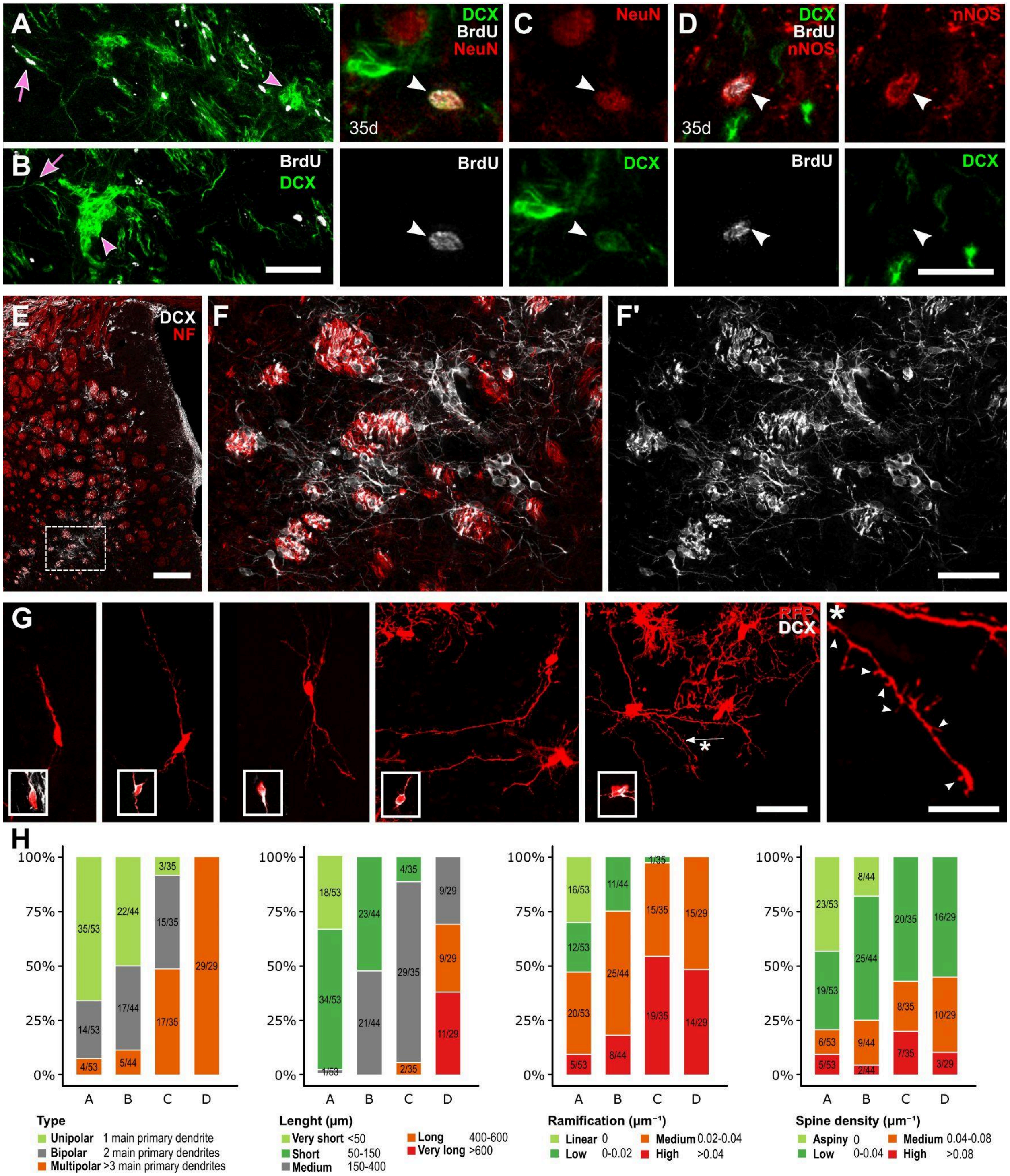


***Supplementary Figure 1. Lesion-induced neuron survival, maturation and morphology.*** **A-B)** Examples of BrdU+/DCX+ (white arrows) and BrdU+/DCX- cells, including both clustered (pink arrowheads) and individual (pink arrows) STR-nbl 15d **(A)** and 35d post BrdU injection **(B)** At both 14

(n=8) and 35 (n=6) days post-BrdU injection almost all BrdU+/DCX+ STR-nbl were already individually dispersed (14d:96±5%, 35d:96±8%). **C)** Representative newborn neuron expressing NeuN that retains DCX expression. **D)** Example of a DCX- newborn neuron expressing nNos. **E-F)** Part of neuroblasts localized within the grey matter, while others resided in the internal capsule fiber bundles, visualized by neurofilament staining (NF, red).(F) Higher-magnification image showing both neuroblast subpopulations. Arrowheads indicate neuroblast residing in the internal capsule fiber bundle. **G)** Confocal image of reconstructed neurons shown in Fig.1. (*) High-magnification of a dendrite (white arrow) with spines indicated by arrowheads. For each neuron, an inset of the cell body shows DCX positivity. **H)** Cluster composition of neuroblasts with different morphological parameters, discretized as indicated, expressed as percentage. Scale bars: 50 µm in A-B G, 20 µm in C-D, 200 µm in E, 75 µm in F.

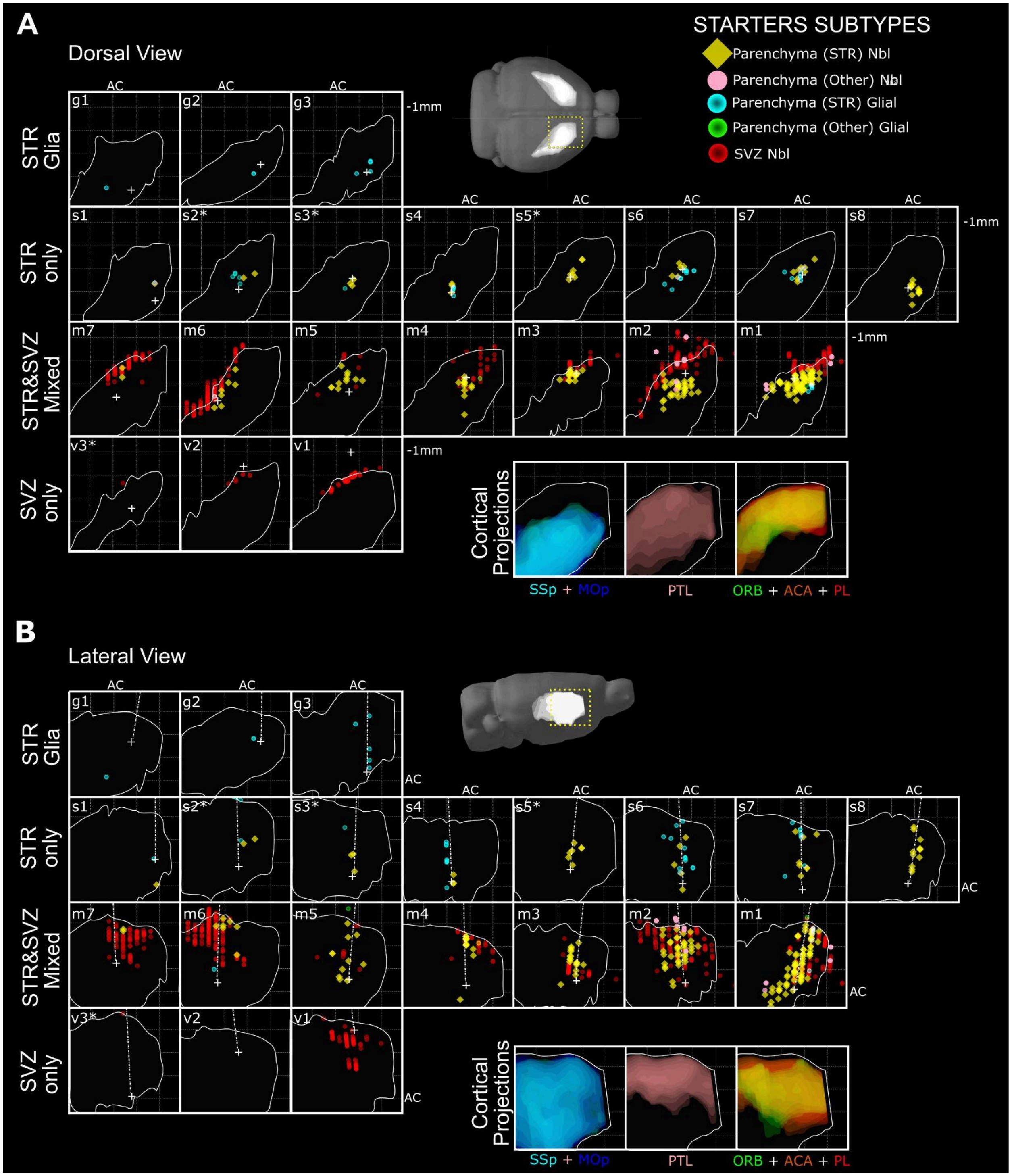


***Supplementary Figure 2. Starter cell distribution in the striatum.*** For each animal, the dorsal **(A)** and lateral views **(B)** are shown. The different starter subtypes are shown with different colors. White crosses in **(A)** and dashed straight lines in **(B)** correspond to the needle track.

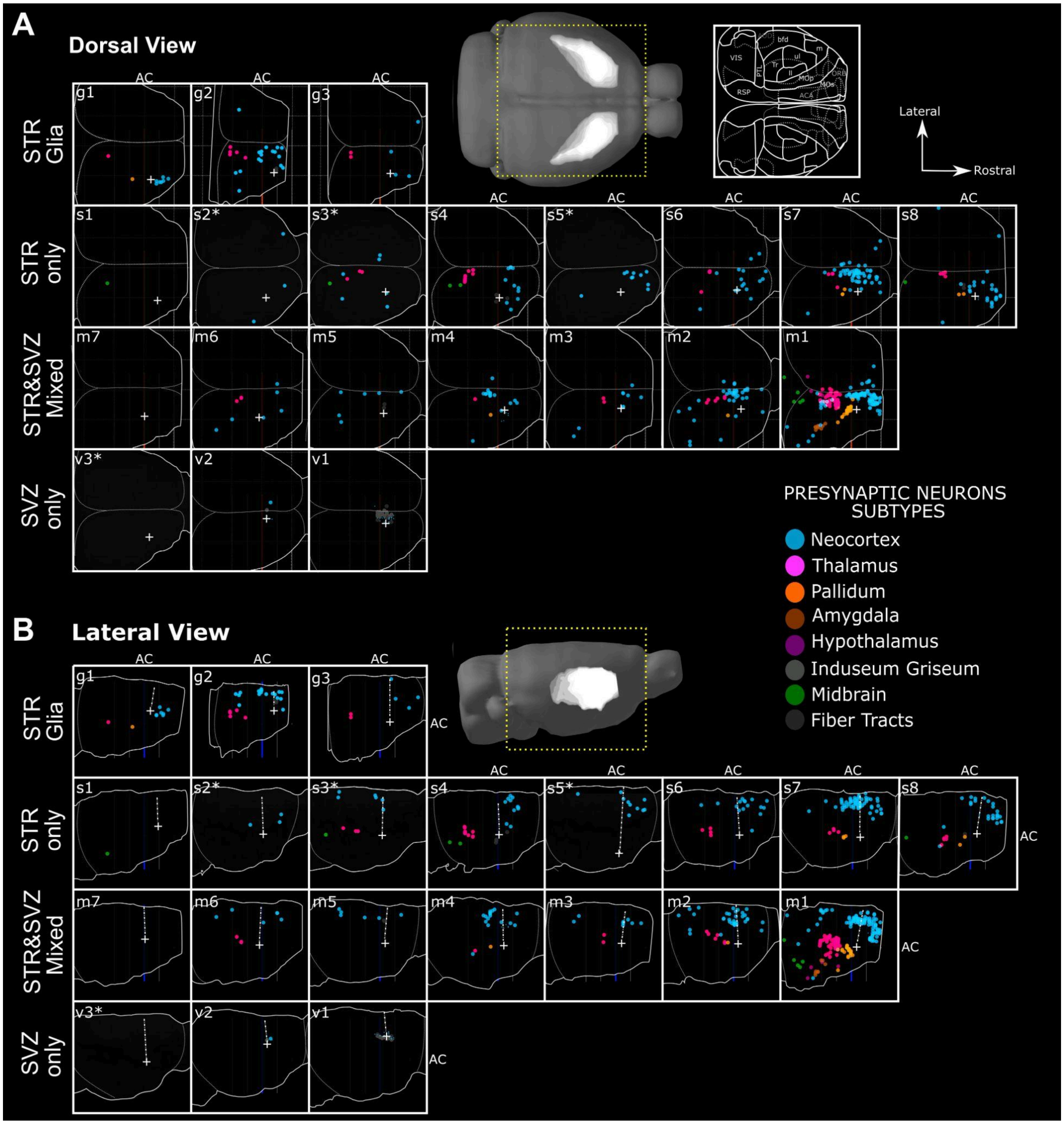


***Supplementary Figure 3. Presynaptic cell distribution.*** For each animal, the dorsal **(A)** and lateral views **(B)** are shown. The different presynaptic cell subtypes are shown with different colors. White crosses in **(A)** and dashed straight lines in **(B)** correspond to the needle track.

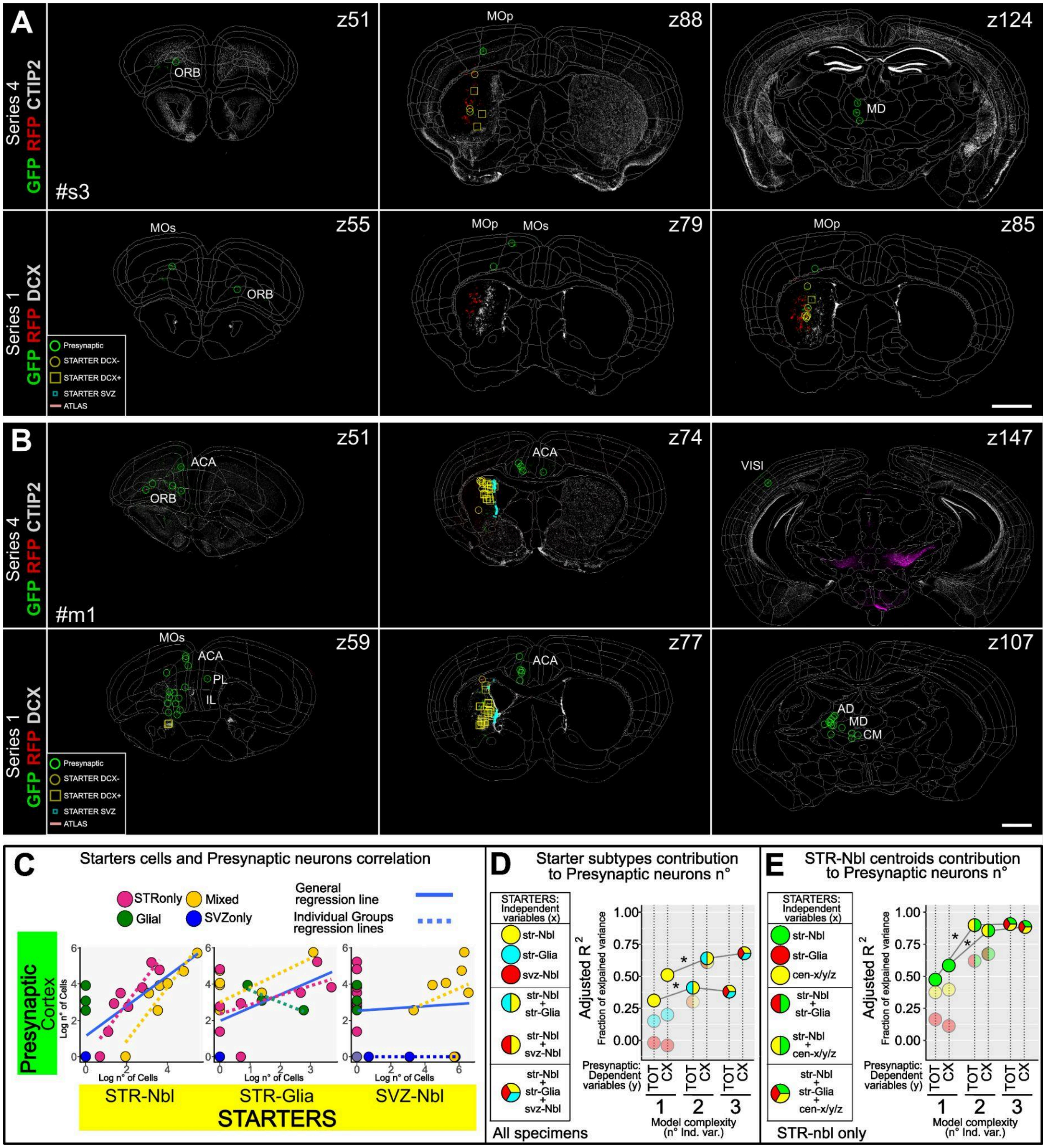

***Supplementary Figure 4. Linear regression analyses of presynaptic neuron distribution.***
**A,B)** Representative sections of two specimens (#s3 in A; #m1 in B) stained for CTIP2 to highlight the striatum and the lesioned area (upper panels) and for DCX to highlight STR-nbl (lower panels). Images were aligned to the Allen Brain Reference atlas. Starter cells and presynaptic neurons were annotated as indicated in the legend. Z147 in (B) was also stained for TH to highlight dopaminergic neurons. **C)** Correlation between the log of presynaptic neurons in the neocortex and the log of STR-nbl, STR-Glia and SVZ-Nbl starter cells. Specimen groups are shown in colour. The regression line for all specimens is in blue, while that of individual specimen groups is dotted and color-coded (see also SupplementaryTable1). **D)** Linear regression analysis evaluating the correlation between the number of different starter subtypes (independent variables) and that of presynaptic neurons either over the entire brain (TOT) or in the neocortex only (CX; dependent variables). The fraction of explained variance in the

number of presynaptic neurons among specimens (adjusted R2; AdjR2) is plotted for different linear models in which the starter subtypes were tested either individually (model complexity=1) or in combination with others (model complexity=2 or 3; see also SupplementaryTable1 for the equations). The dependent variable composition of each model is indicated in the legend. For each level of model complexity, the best models, with higher R2, are in clear color. STR-nbl (yellow) was the best single variable to explain both TOT and CX neurons. The addition of STR-glia (cyan) produced a small but significant increase in Adj.R2 (see also SupplementaryTable1). **E)** Similar to F but testing the contribution of the STR-nbl centroid positions, expressed as x,y,z coordinates (shown in D), to explain the variance in the number of TOT and CX presynaptic neurons (AdjR2). The higher connectivity ratio in STR-only specimens could depend both on glial cells, more abundant in STR-only specimens, and to higher connectivity of the more ventro-lateral STR-nbl of STR-only specimens, possibly by virtue of their larger dendritic arbours. STR-nbl centroids were a better predictor of presynaptic neuron numbers than glial cells, both individually or in combination with STR-nbl, the best individual predictor, in a two-variable model. Only specimens with STR-nbl starters were considered for this analysis. This analysis indicate that STR-nbl contribute much more substantially to observed connectivity than glial cells.

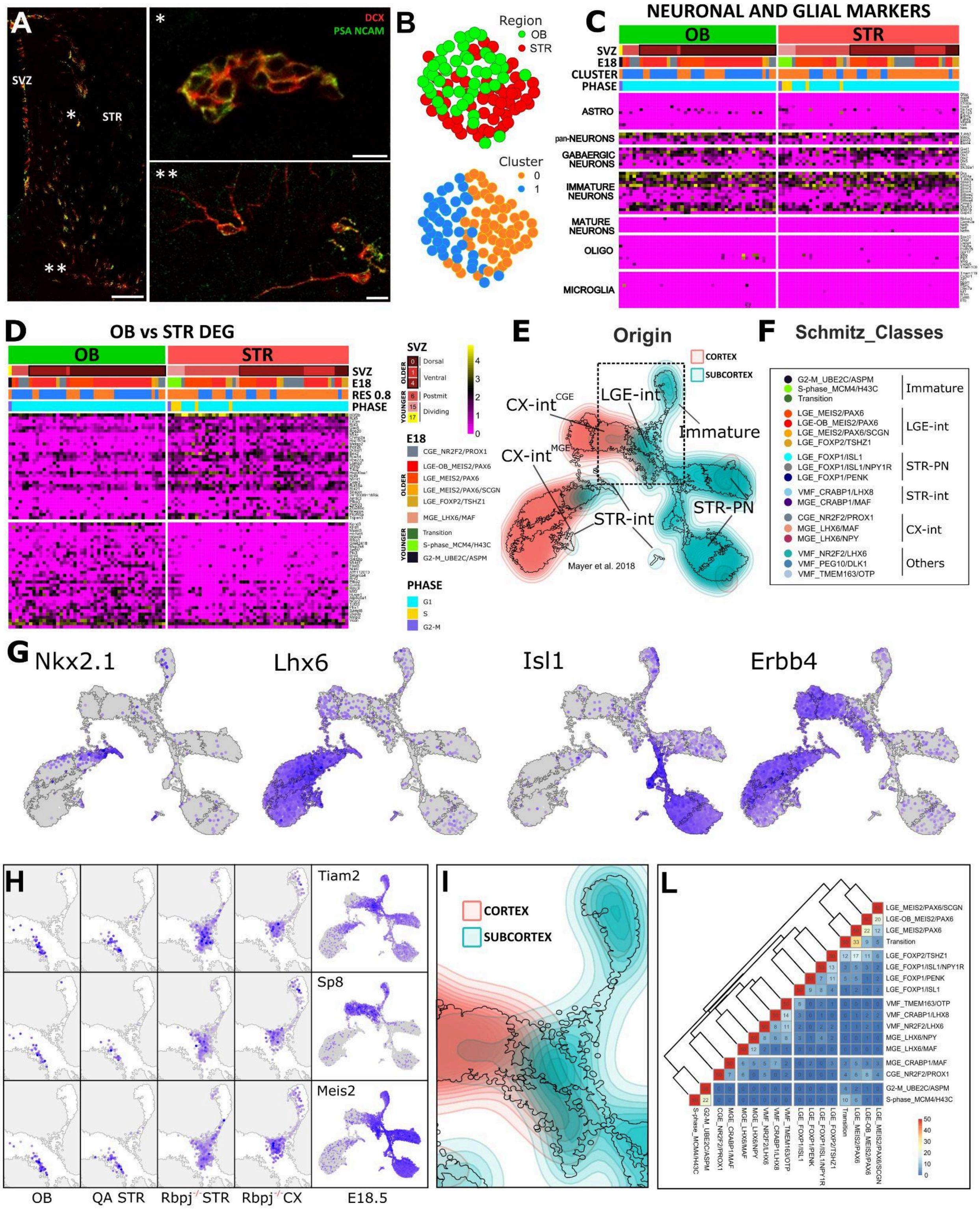

A
DCX
PSA NCAM
SVZ
STR
B
Region
OB
STR
Cluster
0
1
C
NEURONAL AND GLIAL MARKERS
OB
STR
SVZ
E18
CLUSTER
PHASE
ASTRO
pan-NEURONS
GABAERGIC NEURONS
IMMATURE NEURONS
MATURE NEURONS
OLIGO
MICROGLIA
D
OB vs STR DEG
OB
STR
SVZ
E18
RES 0.8
PHASE
SVZ
OLDER
YOUNGER
Dorsal
Ventral
Postmit
Dividing
E18
CGE_NR2F2/PROX1
LGE-OB_MEIS2/PAX6
LGE_MEIS2/PAX6
LGE_MEIS2/PAX6/SCGN
LGE_FOXP2/TSHZ1
MGE_LHX6/MAF
Transition
S-phase_MCM4/H43C
G2-M_UBE2C/ASPM
PHASE
G1
S
G2-M
E
Origin
CORTEX
SUBCORTEX
CX-int^CGE
CX-int^MGE
LGE-int
Immature
STR-int
STR-PN
Mayer et al. 2018
F
Schmitz_Classes
G2-M_UBE2C/ASPM
S-phase_MCM4/H43C
Transition
Immature
LGE_MEIS2/PAX6
LGE-OB_MEIS2/PAX6
LGE_MEIS2/PAX6/SCGN
LGE_FOXP2/TSHZ1
LGE-int
LGE_FOXP1/ISL1
LGE_FOXP1/ISL1/NPY1R
LGE_FOXP1/PENK
STR-PN
VMF_CRABP1/LHX8
MGE_CRABP1/MAF
STR-int
CGE_NR2F2/PROX1
MGE_LHX6/MAF
MGE_LHX6/NPY
CX-int
VMF_NR2F2/LHX6
VMF_PEG10/DLK1
VMF_TMEM163/OTP
Others
G
Nkx2.1
Lhx6
Isl1
Erbb4
H
Tiam2
Sp8
Meis2
OB
QA STR
Rbpj^-/- STR
Rbpj^-/- CX
E18.5
I
CORTEX
SUBCORTEX
L
LGE_MEIS2/PAX6/SCGN
LGE-OB_MEIS2/PAX6
LGE_MEIS2/PAX6
Transition
LGE_FOXP2/TSHZ1
LGE_FOXP1/ISL1/NPY1R
LGE_FOXP1/PENK
LGE_FOXP1/ISL1
VMF_TMEM163/OTP
VMF_CRABP1/LHX8
VMF_NR2F2/LHX6
MGE_LHX6/NPY
MGE_LHX6/MAF
MGE_CRABP1/MAF
CGE_NR2F2/PROX1
G2-M_UBE2C/ASPM
S-phase_MCM4/H43C

***Supplementary Figure 5. Adult neuroblasts sc-RNAseq clustering and E18 projection.***

**A)** Coronal sections of a QA lesioned specimen, 5wpl, at the level of the striatal neurogenic response. Higher magnifications show that DCX+ neuroblasts organized both as clusters (*) and as individual cells (**), and are strongly and specifically labelled by PSA-NCAM. **B)** UMAP space showing clustering of OB and STR cells. Cluster analysis identified two main clusters, which included cells from both regions, with STR cells preferentially in cluster 0 (77%,41/53) while OB cells in cluster 1 (65%, 30/46). **C)** Expression of main neural lineage markers. For neurons, additional markers of mature, immature, and GABAergic neurons are shown. Both OB and STR cells express markers of immature GABAergic neurons but not of mature neurons or glial cells. Cell annotations refer to origin (OB or STR), label transfer on SVZ or Mayer-E18, clustering, and cell cycle phase. Legend as in D. **D)** Top 20 Differentially expressed genes between OB and STR cells. **E)** Density plot showing the overlap of cortical and subcortical-derived cells in Mayer-E18. While cortical MGE and CGE interneurons were mainly isolated from the cortex, and striatal projection neurons from the subcortex, LGE interneurons were distributed in both regions. Note that striatal interneurons also partially overlapped with MGE interneurons, highlighting the strong similarity between these neuron types. **F)** GABAergic neurons classes as defined by Schmitz et al.[9]**. G)** Feature plots showing the expression of main lineage markers in Mayer-E18 cells. Nkx2.1 is expressed by MGE progenitors and striatal interneurons, Lhx6 stains cortical and subcortical MGE-derived interneurons, Isl1 is expressed by D1 MSN, Erbb4 is expressed by all interneurons, derived from LGE, CGE and MGE. Note that positive cells are on top, to highlight the specificity of these gene expression patterns. **H)** Feature plots of the expression of main LGE-MEIS2/PAX6 class interneurons markers in OB, parenchymal neuroblasts (high magnification of LGE interneurons area), and Mayer-E18 cells plotted on Mayer-E18 UMAP space. **I)** Higher magnification of E showing the overlap between cortical and subcortical mature LGE interneurons. **L)** Table of the top 50 genes shared by the main neuron classes in the Mayer-E18 dataset. The 3 LGE MEIS2-PAX6 classes were highly related (20-22 shared markers), but interestingly, the OB and SCGN subtypes shared consistently fewer genes than the other with the transition class that includes very immature, early post-mitotic cells from the subcortical germinative regions (9 and 5 vs 33 / top50 genes shared relative to the LGE MEIS2-PAX6 class). This strongly suggests that these letter classes are at a more advanced maturation stage. Scale bars: 200 µm in B and 10 µm in B*,B**.

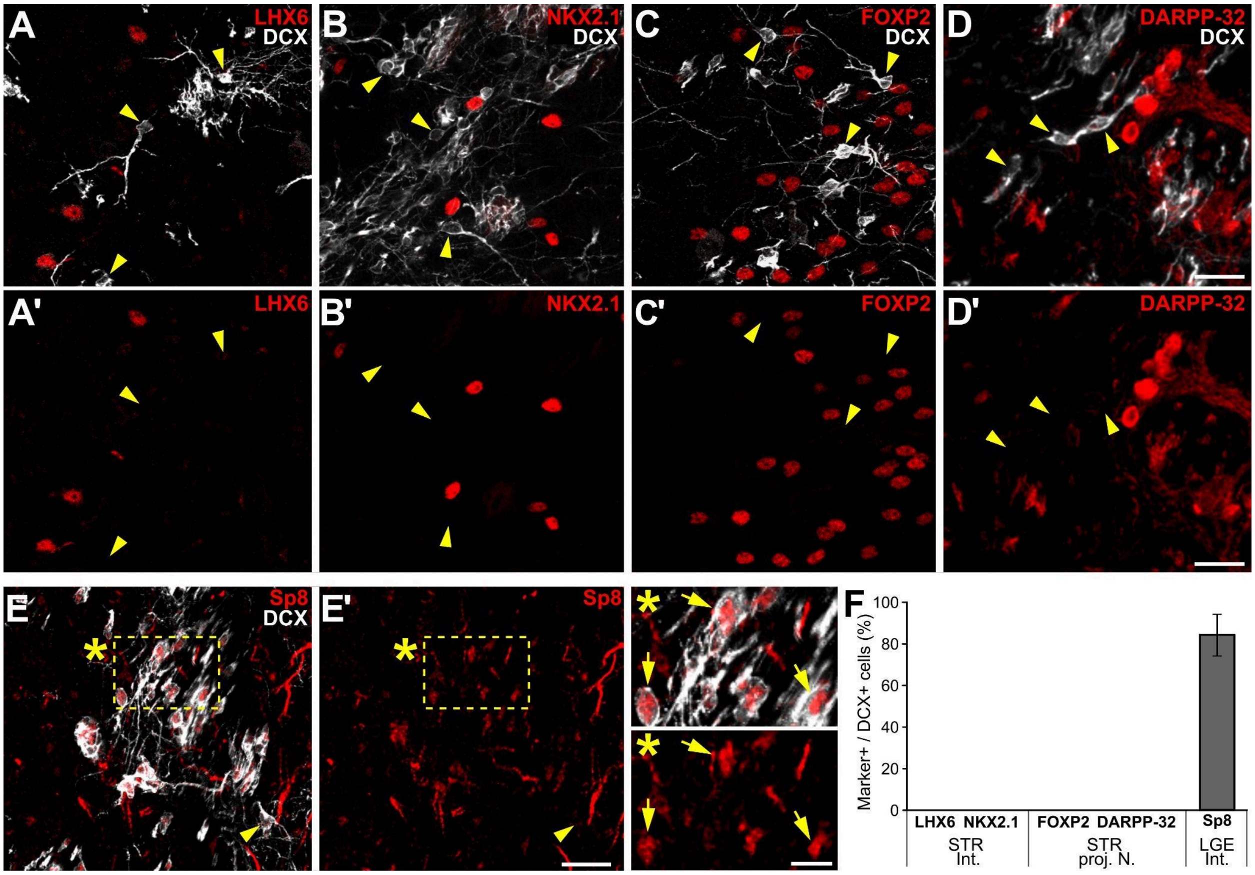


***Supplementary Figure 6. Immunohistochemical phenotyping of QA-lesion induced neuroblasts.***

**A-E')** Representative images from the striatum of specimens at 5 weeks after QA lesion. STR-nbl do not express markers of striatal interneurons (NKX2.1 and LHX6, A-B'), nor of striatal projection neurons (FOXP2 and DARPP-32, C-D'), but express the LGE interneuron marker Sp8 (E,E'). **F)** Quantifications of striatal interneuron, projection neuron, and LGE interneuron markers in DCX+ STR-nbl at 5 wpl (data are presented as mean±SD). Scale bars: 25 µm in A-E', G-I' and 10 µm in E*,E'*.

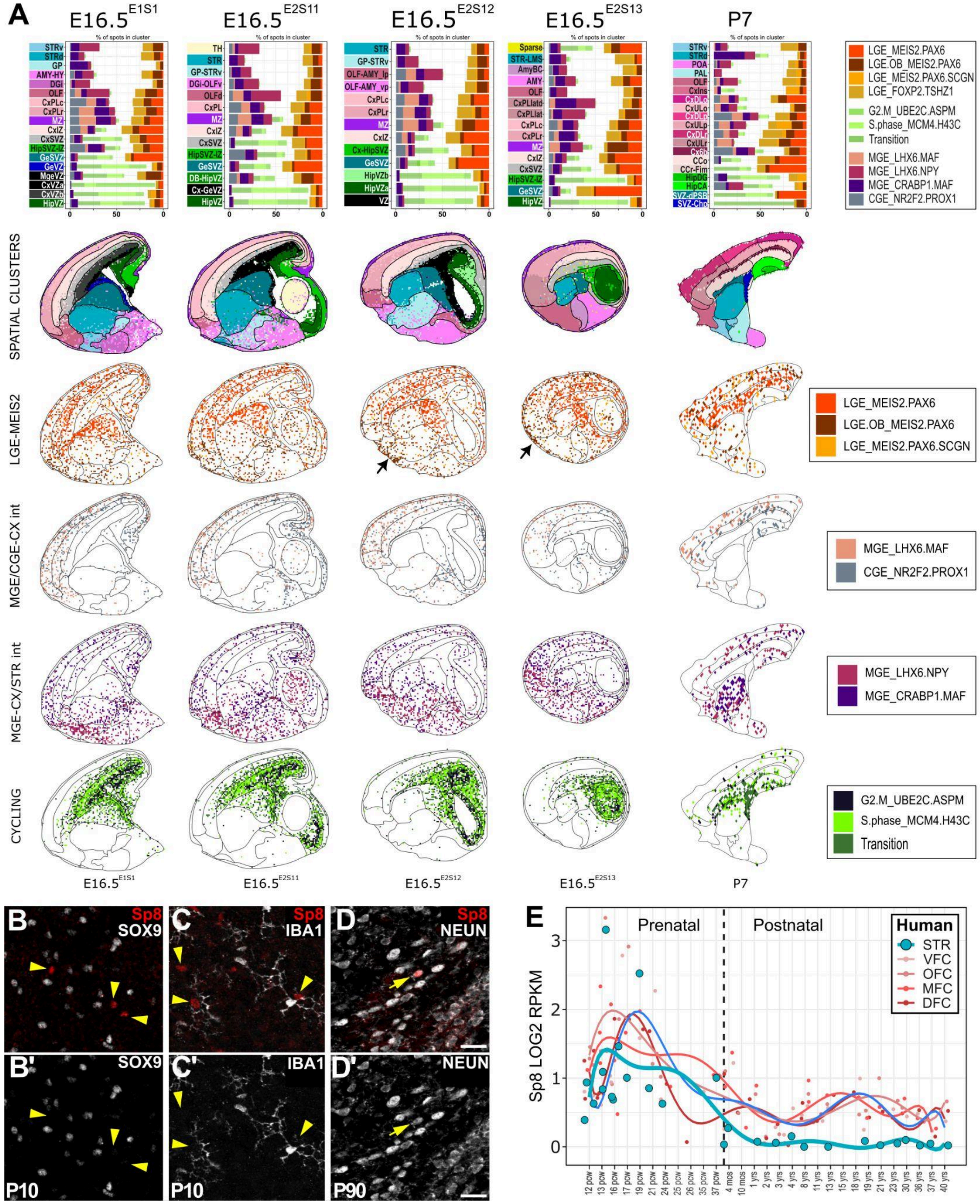


***Supplementary Figure 7. Spatial distribution of LGE_Meis2/Pax6 class cells at E16.5 and P7.***

**A)** Mapping of Mayer-E18 classes onto four E16.5 sagittal sections at different medio-lateral positions and analyzed with stereo-seq and Visium, respectively. The E16.5 sections come from two different specimens (E1 and E2). The counts of the stereo-seq spots, originally of 0.5x0.5µm, were aggregated into bigger areas of 25x25µm. Unbiased clustering has been used to divide spots into areas. Each section has been analyzed separately, and clustering levels were chosen at approximately the same hierarchical level of areal subdivisions. A color code represents each area, and the graphs on top report their relative class composition. Each spot was attributed to the class getting the maximum score among all the Mayer-E18 neuron classes. In the following rows, the spatial distribution of spots assigned to LGE_MEIS2/PAX6 classes, to cortical MGE and CGE derived interneurons, MGE derived striatal interneurons, and cortical and striatal MGE_LHX6/NPY (SST) interneurons, proliferating and early post-mitotic cells. Note that, as expected, the cortical interneurons are preferentially distributed in MZ and CP and do not enter the striatum. By contrast, some of the striatal interneurons also map to the cortex; these subtypes, however, are closely related and accordingly, some cortical cell was assigned a striatal interneuron identity also in Mayer-E18 (Fig.S4E). Interestingly, a group of LGE_MEIS2/PAX6 cells accumulates in the superficial part of the piriform cortex (arrows). This area expresses Eomes and GRM1 (data not shown) and likely corresponds to the caudal part of the LOT, where transient guidepost cells have been previously described. **B-C')** Representative images from P10 mouse striatum, showing that Sp8+ cells do not express markers of astrocyte (SOX9, B,B') nor of microglia (IBA1, C,C'). **D,D')** Representative images from P90 mouse striatum, showing that the few Sp8+ cells that are still observed at this stage express the mature neuron marker NEUN. **E**) Sp8 expression during human development, data from Allen Brain, Human Brain Atlas[121]. In B-D', arrowheads indicate marker-negative cells, while arrows highlight marker-positive cells. Scale bar: 25 µm in B-D'.

### SUPPLEMENTARY TABLES

#### *Supplementary Table 1. Results of all statistical analyses presented in the study.*

* Indicates that a post hoc test was performed, please see below

| Graph | Statistical test | Result | P Value | Post-hoc |
|---|---|---|---|---|
| Fig. 1B (BrdU+ DCX+) | Independent samples t-test | t(12) = 3.6729 | 0.003 | |
| Fig. 1B (BrdU+ DCX-) | Independent samples t-test | t(12) = 0.12838 | 0.900 | |
| Fig. 1C (NeuN+BrdU+) | Independent samples t-test | t(6) = -1.1633 | 0.289 | |
| Fig. 1C (CalrBrdU/BrdU) | Independent samples t-test | t(6) = 0.11058 | 0.916 | |
| Fig. 1C (nNOSBrdU/BrdU) | Independent samples t-test | t(6) = -1.5106 | 0.182 | |
| Related to Fig. 2 (IC vs GM cells) | Independent samples t-test | t(127) = 0.837 | 0.404 | |
| Fig. 3C | One-Way ANOVA | F(2,9)= 9.93 | 0.005 | * |
| Fig. 3D | One-Way ANOVA | F(2,9)= 2.18 | 0.169 | |
| Fig. 5C (total length) | Generalized Linear Mixed Effects Model (Neg. Binom.) | β = 0.66, SE = 0.22, z = 2.99 | 0.022 | |
| Fig. 5C (branching points) | Generalized Linear Mixed Effects Model (Poisson) | β = 0.99, SE = 0.28, z = 2.99 | 0.013 | |
| Fig. 5D (ML) | Generalized Linear Mixed Effects Model (Tweedie) | β = -3.03, SE = 0.74, z = -4.12 | 0.0002 | |
| Fig. 5D (DV) | Generalized Linear Mixed Effects Model (Tweedie) | β = -0.57, SE = 0.18, z = -3.09 | 0.002 | |
| Related to Fig. 5D (AP) | Linear Mixed Effects Model | β = 0.06, SE = 0.09, t = 0.65 | 0.524 | |
| Fig. 5E (TOTAL) | Welch Two Sample t-test | t(7.62)= -3.04 | 0.017 | |
| Fig. 5E (CORTEX total) | Welch Two Sample t-test | t(8.12)= -4.01 | 0.004 | |
| Fig. 5E (CORTEX ipsi) | Welch Two Sample t-test | t(7.88)= -3.63 | 0.007 | |
| Fig. 5E (CORTEX contra) | Welch Two Sample t-test | t(7.07)= -2.25 | 0.059 | |
| Fig. 5E (THALAMUS) | Welch Two Sample t-test | t(7.39)= -1.34 | 0.221 | |
| Fig. 5E (PALLIDUM) | Welch Two Sample t-test | t(12.5)= 0.37 | 0.715 | |
| Fig. 5E (MIDBRAIN) | Welch Two Sample t-test | t(7.01)= -1.75 | 0.123 | |
| Fig. S3C | Multiple Linear Regression | See below | | |
| Fig. S3D | Multiple Linear Regression | See below | | |
| Fig. S3E | Multiple Linear Regression | See below | | |
| Fig. 6E (OB vs STR) | Fisher's Exact Test (only mature cells included; LGE_MEIS2_PAX6 classes pooled together) | | 0.352 | * |
| Fig. 6E (OB vs Rbpj-STR) | Fisher's Exact Test (only mature cells included; LGE_MEIS2_PAX6 classes pooled together) | | 0.012 | * |
| Fig. 6E (OB vs Rbpj-CX) | Fisher's Exact Test (only mature cells included; LGE_MEIS2_PAX6 classes pooled together) | | 0.172 | * |
| Fig. 6E (OB vs STR) | Fisher's Exact Test | | 0.402 | * |

| | | | | |
|---|---|---|---|---|
| | (only mature cells included; LGE_MEIS2_PAX6 classes splitted) | | | |
| Fig. 6E (OB vs Rbpj-STR) | Fisher’s Exact Test (only mature cells included; LGE_MEIS2_PAX6 classes splitted) | | < 2.2e-16 | * |
| Fig. 6E (OB vs Rbpj-CX) | Fisher’s Exact Test (only mature cells included; LGE_MEIS2_PAX6 classes splitted) | | < 2.2e-16 | * |
| Fig. 7H | Independent samples t-test | t(4) = 3.3709 | 0.028 | |
| Fig. 7I | Welch Two Sample t-test | t(3.002) = -3.676 | 0.035 | |
| Fig. S7B (OB vs STR) | Fisher’s Exact Test (Mature and Immature clusters pooled together) | | 0.002 | |
| Fig. S7B (OB vs Rbpj-STR) | Fisher’s Exact Test (Mature and Immature clusters pooled together) | | < 2.2e-16 | |
| Fig. S7B (OB vs Rbpj-CX) | Fisher’s Exact Test (Mature and Immature clusters pooled together) | | < 2.2e-16 | |
| Fig. S7B (STR vs Rbpj-STR) | Fisher’s Exact Test (Only Immature clusters) | | 0.374 | |
| Fig. S7B (STR vs Rbpj-CX) | Fisher’s Exact Test (Only Immature clusters) | | 0.185 | |
| Fig. S7B (OB vs STR) | Fisher’s Exact Test (Only Mature clusters) | | 5.1e-07 | * |
| Fig. S7B (OB vs Rbpj-STR) | Fisher’s Exact Test (Only Mature clusters) | | 0.260 | * |
| Fig. S7B (OB vs Rbpj-CX) | Fisher’s Exact Test (Only Mature clusters) | | 0.024 | * |
| Fig. S7B (STR vs Rbpj-STR) | Fisher’s Exact Test (Only Mature clusters) | | 6.4e-15 | * |
| Fig. S7B (STR vs Rbpj-CX) | Fisher’s Exact Test (Only Mature clusters) | | 9.5e-10 | * |

### Post hoc analyses

| **Figure 3C. Post hoc** | |
|---|---|
| Comparison | Tukey |
| Type1 - Type2 | 0.072 |
| Type1 - Type3 | 0.004 |
| Type2 - Type3 | 0.068 |

| **Figure 6E. Pairwise Fisher’s test** | | | |
|---|---|---|---|
| **Comparison (LGE_MEIS2_PAX6 classes pooled together)** | **OB vs STR** | **OB vs Rbpj-STR** | **OB vs Rbpj-CX** |
| CGE_NR2F2.PROX1 | 1 | 0.014 | 1 |
| LGE_MEIS2.PAX6.ALL | 0.542 | 0.014 | 0.710 |
| LGE_FOXP2.TSHZ1 | 0.542 | 0.291 | 0.314 |
| LGE_FOXP1.ISL1 | nd | 1 | nd |

| | | | |
|---|---|---|---|
| MGE_LHX6.MAF | 1 | nd | nd |
| **Comparison (LGE_MEIS2_PAX6 classes splitted)** | **OB vs STR** | **OB vs Rbpj-STR** | **OB vs Rbpj-CX** |
| CGE_NR2F2.PROX1 | 1 | 0.014 | 1 |
| LGE.OB_MEIS2.PAX6 | 0587 | < 0.001 | < 0.001 |
| LGE_MEIS2.PAX6 | 0.954 | < 0.001 | < 0.001 |
| LGE_MEIS2.PAX6.SCGN | 1 | 0.147 | 0.408 |
| LGE_FOXP2.TSHZ1 | 0.813 | 0.262 | 0.175 |
| LGE_FOXP1.ISL1 | nd | 1 | nd |
| MGE_LHX6.MAF | 1 | nd | nd |

| **Figure S7B. Pairwise Fisher's test** | | | | | |
|---|---|---|---|---|---|
| **Comparison** | **OB vs STR** | **OB vs Rbpj-STR** | **OB vs Rbpj-CX** | **STR vs Rbpj-STR** | **STR vs Rbpj-CX** |
| Cluster 0 | < 0.001 | 0.417 | 0.034 | < 0.001 | < 0.001 |
| Cluster 1 | 0.006 | 0.456 | 1 | < 0.001 | 0.019 |
| Cluster 4 | 0.009 | 0.417 | 0.035 | < 0.001 | < 0.001 |

### Figures S3C,D

| | | **MODEL** | | **TOT** | **CX** | **TH** |
|---|---|---|---|---|---|---|
| **Fig. S3 C** | **1.1** | PreS= $\beta_0 + \beta_1$STRnbl + ε | **p** | 0,0040 | 0,0001 | 0,0054 |
| | | | **Adj R2** | **0,31** | **0,51** | **0,29** |
| | **1.2** | PreS= $\beta_0 + \beta_1$STRglia + ε | **p** | 0,0408 | 0,0215 | 0,0008 |
| | | | **Adj R2** | **0,15** | **0,20** | **0,41** |
| | **1.3** | PreS= $\beta_0 + \beta_1$SVZ + ε | **p** | 0,4293 | 0,6948 | 0,5085 |
| | | | **Adj R2** | -0,02 | -0,04 | -0,03 |

**Fig. S3D**

| | | | | | | |
|---|---|---|---|---|---|---|
| **2 Variables** | **2.1** | PreS= $\beta_0 + \beta_1$STRnbl + $\beta_2$STRglia + ε | **p** | 0,0025 | 0,0000 | 0,0000 |
| | | | **Adj R2** | **0,41** | **0,64** | **0,62** |
| | **2.2** | PreS= $\beta_0 + \beta_1$STRnbl + $\beta_2$SVZ + ε | **p** | 0,0128 | 0,0001 | 0,0154 |
| | | | **Adj R2** | **0,30** | **0,61** | **0,29** |
| **3 Var** | **3.1** | PreS= $\beta_0 + \beta_1$STRnbl + $\beta_2$STRglia + $\beta_3$SVZ + ε | **p** | 0,0080 | 0,0308 | 0,0002 |
| | | | **Adj R2** | **0,38** | **0,68** | **0,60** |

| **MODEL COMPARISONS (PARTIAL F-TEST)** | | | | |
|---|---|---|---|---|
| **1.1** (STRnbl) vs **3.1** (All starters) | **p** | 0,153 | 0,010 | 0,052 |
| **1.1** (STRnbl) **vs** **2.1** (STRnbl + STRglia) | **p** | 0,050 | 0,010 | 0,000 |
| **1.1** (STRnbl) **vs** **2.2** (STRnbl + SVZ) | **p** | 0,432 | 0,024 | 0,368 |

**Fig.S3E**

| | | MODEL | | TOT | CX | TH |
|---|---|---|---|---|---|---|
| 1 Variable | 1.1 | PreS= $\beta_0 + \beta_1$STRnbl + ε | p | 0,0025 | 0,0005 | 0,0182 |
| | | | Adj R2 | **0,48** | **0,59** | **0,31** |
| | 1.2 | PreS= $\beta_0 + \beta_1$STRglia + ε | p | 0,0700 | 0,1124 | 0,0094 |
| | | | Adj R2 | **0,17** | **0,12** | **0,37** |
| | 1.3 | PreS= $\beta_0 + \beta_1$(X*Y*Z) + ε | p | 0,1547 | 0,1441 | 0,7555 |
| | | | Adj R2 | **0,38** | **0,40** | **-0,27** |
| 2 Variables | 2.1 | PreS= $\beta_0 + \beta_1$STRnbl + $\beta_2$STRglia+ ε | p | 0,0013 | 0,0004 | 0,0009 |
| | | | Adj R2 | **0,62** | **0,68** | **0,64** |
| | 2.2 | PreS= $\beta_0 + \beta_1$STRnbl + $\beta_2$(X*Y*Z)+ ε | p | 0,0016 | 0,0040 | 0,0245 |
| | | | Adj R2 | **0,90** | **0,86** | **0,73** |
| 3 Var.s | 3.1 | PreS= $\beta_0 + \beta_1$STRnbl + $\beta_2$STRglia + $\beta_3$(X*Y*Z) + ε | p | 0,0032 | 0,0051 | 0,0246 |
| | | | Adj R2 | **0,91** | **0,89** | **0,79** |

| MODEL COMPARISONS (PARTIAL F-TEST) | | | | | |
|---|---|---|---|---|---|
| 3 vs 2.1 | (Centroids contribution to full model) | p | 0,027 | 0,064 | 0,199 |
| 3 vs 2.2 | (STR_glia contribution to full model) | p | 0,218 | 0,156 | 0,162 |
| 2.2 vs 1.1 | (Centroids addition to Nbl) | p | 0,010 | 0,043 | 0,061 |
| 2.2 vs 1.1 | (Glia addition to Nbl) | p | 0,036 | 0,050 | 0,004 |
| 2.2 vs 1.1 | (Centroids addition vs Glia addition to Nbl) | p | 0,020 | 0,077 | 0,275 |

**COORDINATE CONTRIBUTION TO CONNECTIVITY RATIO (only Specimens with STR-nbl)**

| MODEL | | TOT | CX |
|---|---|---|---|
| 3) X*Y*Z | p | 0,0048 | 0,1228 |
| | Adj R2 | **0,80** | **0,43** |
| 2.1) X*Z | p | 0,1877 | 0,3274 |
| | Adj R2 | **0,16** | **0,06** |
| 2.1) X*Y | p | 0,0490 | 0,0597 |
| | Adj R2 | **0,36** | **0,33** |
| 2.2) Y*Z | p | 0,0001 | 0,0416 |
| | Adj R2 | **0,79** | **0,38** |
| 1.1) X | p | 0,1858 | 0,1740 |
| | Adj R2 | **0,06** | **0,07** |
| 1.2) Z | p | 0,0270 | 0,1535 |
| | Adj R2 | **0,27** | **0,08** |
| 1.3) Y | p | 0,0353 | 0,0916 |
| | Adj R2 | **0,24** | **0,14** |

| MODEL COMPARISONS (PARTIAL F-TEST) | | | |
|---|---|---|---|
| X vx X*Z | p | 0,044 | 0,431 |
| X vs X*Y | p | 0,184 | 0,064 |
| X*Z vs X*Y*Z | p | 0,052 | 0,110 |
| X*Y vx X*Y*Z | p | 0,023 | 0,306 |
| Z*Y vs X*Y*Z | p | 0,043 | 0,371 |

***Supplementary Table 2. Stereotaxic coordinates used for monosynaptic retrograde tracing experiments.***

Retro and Rabies viruses were injected at the same coordinates in each specimen. EXP: experimental batch; ID: specimen ID; ML: mediolateral; AP: antero-posterior; DV: dorso-ventral.

| | TARGET STR | | | | | | | | | | | | | | | | | | | TARGET SVZ | | |
|---|---|---|---|---|---|---|---|---|---|---|---|---|---|---|---|---|---|---|---|---|---|---|
| Exp | 2 | 1 | 1 | 2 | 2 | 3 | 3 | 1 | 3 | 1 | 2 | 2 | 1 | 1 | 1 | 1 | 2 | 2 | 3 | 2 | 2 | 2 |
| ID | s8 | s5 | g1 | g2 | g3 | s4 | s6 | m1 | s7 | m2 | s2 | m5 | m3 | s1 | m7 | m6 | m4 | s3 | v3 | x1 | v1 | v2 |
| ML | -2 | | | -1.9 | | | | -1.8 | | | | | | | | | -1.7 | | | -1 | -0.8 | -0.7 |
| AP | 0.8 | | 0.6 | 1.1 | 1 | 0.9 | | 0.9 | | 0.8 | | | | 0.6 | | | 1 | 0.9 | | 1 | 0.6 | |
| DV (-) | 3.3 | 3 | 3 / 3.2 | 3.2 | 3.6 | 3 / 3.4 | 3 / 3.4 | 3.4 | 3.2 / 3.4 | 3.5 | 3.2 / 3.4 | 3.4 | 3 / 3.2 | 3.2 | 3.2 | 3 | 3.5 | 3.5 | 3 / 3.4 | 2 / 2.4 | 2 / 2.4 | 2 / 2.4 |

*Supplementary Table 3. List of primary and secondary antibodies.*

| Antigen Name | Host | Dilution | Source |
|---|---|---|---|
| Primary antibodies | | | |
| BrdU | Rat | 1:1500 | AbD Serotec, Kidlington, UK |
| DCX | Goat | 1:1500 | Santa Cruz Biotechnology, Santa Cruz, CA |
| DCX | Guinea Pig | 1:1500 | Millipore, Bellerica, MA |
| Ki67 | Rabbit | 1:1000 | Abcam, Cambridge, UK |
| NeuN | Mouse | 1:1000 | Chemicon, Temecula, CA |
| DARPP32 | Mouse | 1:1000 | BD Transduction Laboratories, Lexington, KY |
| Calretinin | Rabbit | 1:2000 | Swant, Bellinzona, CH |
| nNOS | Rabbit | 1:6000 | Abcam, Cambridge, UK |
| GFP | Chicken | 1:1000 | Aveslab, Tigard, OR |
| RFP | Rabbit | 1:1000 | Rockland, Philadelphia, PA |
| CTIP2 | Rat | 1:750 | Abcam, Cambridge, UK |
| Sp8 | Rabbit | 1:8000 | Millipore, Bellerica, MA |
| NKX2.1 | Rabbit | Verifica | Biopat s.r.l., Piedimonte Matese, IT |
| LHX6 | Rabbit | Verifica | (Grigoriou et al., 1998) |
| FOXP2 | Rabbit | 1:1000 | Abcam, Cambridge, UK |
| SOX9 | Goat | 1:1200 | R&D Systems, Minneapolis, MN |
| SOX10 | Goat | 1:750 | Santa Cruz Biotechnology, Santa Cruz, CA |
| IBA1 | | | FUJIFILM Wako, Osaka, JP |
| Secondary antibodies | | | |
| Cy3 Anti-Rb,Ms,Gt,Rt,Gp | Donkey | 1:800 | Jackson ImmunoResearch, West Grove, PA |
| Alexa488 Anti-Rb,Ms,Gt,Rt,Ck | Donkey | 1:400 | Jackson ImmunoResearch, West Grove, PA |
| Alexa647Anti-Rb,Ms,Gt,Rt | Donkey | 1:800 | Jackson ImmunoResearch, West Grove, PA |
| Alexa594 Anti-Gt | Donkey | 1:400 | Jackson ImmunoResearch, West Grove, PA |
| Byotinilated Anti-Gt,Gp | Horse | 1:100 | VectorLabs, Newark, CA |
| AMCA-avidinD | | 1:100 | VectorLabs, Newark, CA |
